# Human Immunodeficiency Virus 1 Capsid Uncoating in the Nucleus Progresses Through Defect Formation in the Capsid Lattice

**DOI:** 10.1101/2023.08.22.553958

**Authors:** Levi B. Gifford, Gregory B. Melikyan

## Abstract

The HIV-1 core consists of a cone-shaped capsid shell made of ∼250 capsid protein (CA) hexamers and 12 pentamers encapsulating the viral genome. HIV-1 capsid disassembly, referred to as uncoating, is a highly regulated process that is important for productive infection, however, the location, timing, and regulation of uncoating remain controversial. Here, we employ amber codon suppression to directly label CA and visualize capsid trafficking and uncoating in live cells. In addition to direct CA labeling, a fluid phase fluorescent probe is incorporated into the viral core to detect the formation of small defects in the capsid lattice. This double-labeling strategy does not significantly impact HIV-1 infectivity, maturation, nuclear import, or capsid stability. Single virus tracking reveals nuclear import of intact cores defined as complexes containing both the fluid phase marker and robust CA signal. Subsequent uncoating of HIV-1 cores in the nucleus is manifested by a sequential loss of both fluorescent markers. This two-step uncoating – release of the core content marker followed by loss of CA – is observed in different cells, including a macrophage line. Importantly, the lag between the two steps of uncoating (∼30 min) appears independent of the cell type and is much longer than upon uncoating of cell-free viruses. These data suggest that HIV- 1 uncoating in the nucleus is initiated through a localized defect in the capsid lattice that precedes a global loss of CA. Our results imply that intact HIV-1 cores enter the cell nucleus and uncoat in a stepwise fashion, before integrating into the host genome.

---

The HIV-1 capsid is a large (60 nm wide end, 40 nm narrow end, ∼100 nm length) proteinaceous structure that is comprised of ∼250 capsid protein (CA) hexamers and exactly 12 pentamers to form the conical capsid lattice.^1–4^ Fusion of HIV-1 with the cell membrane releases the capsid into the cytosol where it interacts with a multitude of cellular dependency and restriction factors. Interactions with host dependency factors promote microtubule transport, import through the nuclear pore complex (NPC), and translocation to nuclear speckles for integration within the speckle-associated genomic domains.^3, 5–9^ Capsid disassembly, referred to as uncoating, is required for the release of the HIV-1 pre-integration complex, but the extent and cellular sites of CA loss remain controversial.^1, 3, 10^ HIV-1 capsid stability is tightly regulated by multiple host factors, such as IP6, Sec24C, Nup153, and several molecular motors.^11–14^ Optimal core stability is essential for nuclear import and delivery of viral complexes to the sites of integration, as evidenced by the compromised infectivity of HIV-1 containing CA mutations that hyper-stabilize or destabilize the capsid lattice.^2, 15–18^ Thus, timely HIV-1 uncoating is critical for productive infection.

HIV-1 uncoating has been traditionally studied using biochemical and functional assays that yielded discrepant findings regarding the sites and timing of capsid disassembly.^19–23^ More recently, single HIV-1 imaging in living cells has been employed to visualize the sites and timing of uncoating in the context of productive infection.^6, 24–27^ However, contradicting results regarding the sites of productive uncoating have been reported using single virus imaging approaches in live and fixed cells.^6, 24–34^ Models proposed based upon imaging, biochemical and functional experiments place HIV-1 uncoating in all three possible cellular compartments: the cytosol, shortly after viral fusion,^19–21, 24, 35^, the nuclear pore complex,^28–30, 32, 36^ and the nucleoplasm,^25, 27, 33, 34, 36–39^ with terminal disassembly near the sites of integration.^25, 27^

Recent correlative light and electron microscopy (CLEM) and electron tomography (ET) experiments have detected cracked capsid lattices and, in rare cases, intact-looking capsids within the nucleus of cells.^33, 38^ Although a link between seemingly intact capsid cores in the nucleus and infection could not be established, this finding implies that cores may not lose the capsid lattice upon translocation through the NPC. These CLEM results are consistent with the recent finding that the diameter of intact nuclear pore is not ∼40 nm, as was previously thought, but 60-64 nm, which can, in principle, allow for passage of the entire capsid into the nucleus.^38, 40^ These results are also in line with biochemical and genetic evidence suggesting that, at least a fraction of capsid lattice, is preserved in the nucleus^41^ and that viral cores undergo remodeling upon nuclear import.^42–44^ The presence of cracked cores, which lack the electron density associated with the viral RNP (vRNP), in the nucleoplasm has been taken to indicate that uncoating does not culminate in full disassembly of capsid lattice.^33, 38^

The controversial findings regarding the cellular sites and mechanism of HIV-1 capsid uncoating obtained by single virus imaging approaches appear due, in a large part, to the lack of a minimally invasive direct CA labeling scheme. Because the HIV-1 capsid is a large and highly complex structure that interacts with multiple host dependency factors and plays critical roles in early infection,^2, 3, 39, 45^ most mutations disrupt the capsid morphology, stability and/or reduce infectivity (e.g., ^2, 46^). For this reason, several studies relied on indirect capsid markers to investigate capsid uncoating. One indirect labeling approach utilizes a fluid phase GFP content marker derived from the HIV-1 Gag construct with "internal” GFP (iGFP).^24, 27, 47, 48^ Release of a small portion of iGFP trapped in intact HIV-1 cores has been used as a proxy for uncoating.^24, 27, 48^ However, single particle imaging experiments using this core content marker by two groups led to contradicting conclusions regarding the sites of HIV-1 uncoating.^24, 27^ Another indirect HIV-1 capsid marker, cyclophilin A-DsRed (CDR), utilized cyclophilin A that is rendered tetrameric through fusion to DsRed protein and that tightly binds to the cyclophilin A binding loops of capsid.^6, 26, 30, 49^ We have previously observed loss of this marker after virus docking at the NPC, prior to nuclear import and productive infection.^6, 26, 30, 49^

Direct capsid labeling approaches include a tetracysteine-tagged^32, 50^ or eGFP-tagged CA.^25, 32, 44^ However, both approaches compromise the infectivity and require a large (10-15-fold) excess of WT CA to produce functional virions. Additionally, non-specific labeling and significant photobleaching of the tetracysteine tag during live cell imaging limit its utility.^32, 50^ By tracking HIV-1 cores containing the eGFP-CA probe, Burdick and co-authors have observed nuclear import of HIV-1 core without detectable loss of eGFP-CA, followed by loss of the capsid marker in the nucleus, near the site of integration.^25^ More recently, amber codon suppression^51–55^ has been utilized to label the capsid with non-canonical amino acids conjugated to organic fluorophores.^34^ Insertion of a non-canonical amino acid at the alanine-14 position of CA produced infectious viruses, but grossly delayed nuclear import of HIV-1 complexes, suggesting altered interactions with the nuclear pore complex and/or other host dependency factors. Comparison of labeled capsid signals in the nuclei of fixed cells and to cell-free labeled cores on coverslips revealed no differences, further supporting the notion that intact cores can enter the nucleus.^34^

To investigate the site(s) and dynamics of capsid uncoating, we sought to implement a minimally invasive capsid labeling approach. Toward this goal, we utilized amber codon suppression to label CA at the isoleucine 91 site within the cyclophilin A binding loop. When mixed with comparable amounts of wild-type (WT) CA, the I91 CA mutant containing virions exhibit infectivity close to that of unlabeled pseudoviruses, while allowing for efficient direct labeling of viral capsid and tracking single core uncoating events that result in productive infection. Along with amber codon suppression-based labeling, we produced dual-labeled HIV-1 pseudoviruses containing the fluid phase viral content marker Gag-iYFP (similar to Gag-iGFP^24, 27, 47, 48^) to visualize the relationship between the initial loss of lattice integrity and terminal disassembly of single capsid cores in live cells. With this system, we show that, both the initial loss of integrity and terminal disassembly of capsid leading to productive infection, occur in the nucleus, in agreement with the recent model for nuclear uncoating.^25, 27^ We also find that terminal capsid disassembly is temporally linked to loss of lattice integrity, where the formation of initial/small defect(s) eventually culminates in capsid uncoating. The two-step uncoating phenotype is observed in both osteosarcoma cells and THP-1 macrophage line. Our results provide important insights into the dynamic process of capsid uncoating that progresses through at least one distinct intermediate step preceding terminal loss of CA. These findings are essential for understanding the mechanisms of regulation of the HIV-1’s capsid stability which is critical for productive infection.

## Results

### Characterization and validation of direct HIV-1 capsid protein labeling with unnatural amino acids

To directly label the HIV-1 capsid, we genetically tag the CA protein at various amino acid sites using amber codon suppression. Based on previous study describing CA labeling with amber codon suppression^34^ and analysis of published CA hexamer structures,^56, 57^, we chose four sites to label the CA lattice. From the four amino acid sites, we selected the isoleucine-91 (I91) site within the cyclophilin A binding loop, which allowed direct virus labeling (see Methods for virus production and labeling protocols), while not considerably impairing HIV-1 infectivity or the kinetics of nuclear import, in contrast to the other CA mutants chosen. Incorporation of the non-canonical amino acid trans-cyclooctene lysine (TCO)^51, 52^ in the context of ΔEnv HIV-1 backbone allowed strain-promoted inverse electron-demand Diels-Alder cycloaddition click-labeling^34, 52, 53, 55, 58^ of CA with tetrazine-conjugated fluorophores (referred to as CA*, Fig. 1A). We found that the infectivity of pseudoviruses made of pure CA* and click-labeled with organic fluorophores at either of the four chosen CA positions is significantly diminished. To produce infectious CA* virus, virus-producing cells were transfected at a 1.5:1 ratio of respective plasmids provided. Neither Gag cleavage/maturation nor specific infectivity of pseudoviruses with mixed CA*/CA^WT^ cores labeled at the CA position I91 with SiR-tetrazine was majorly impacted (Fig. 1B, C).

**Fig 1.**
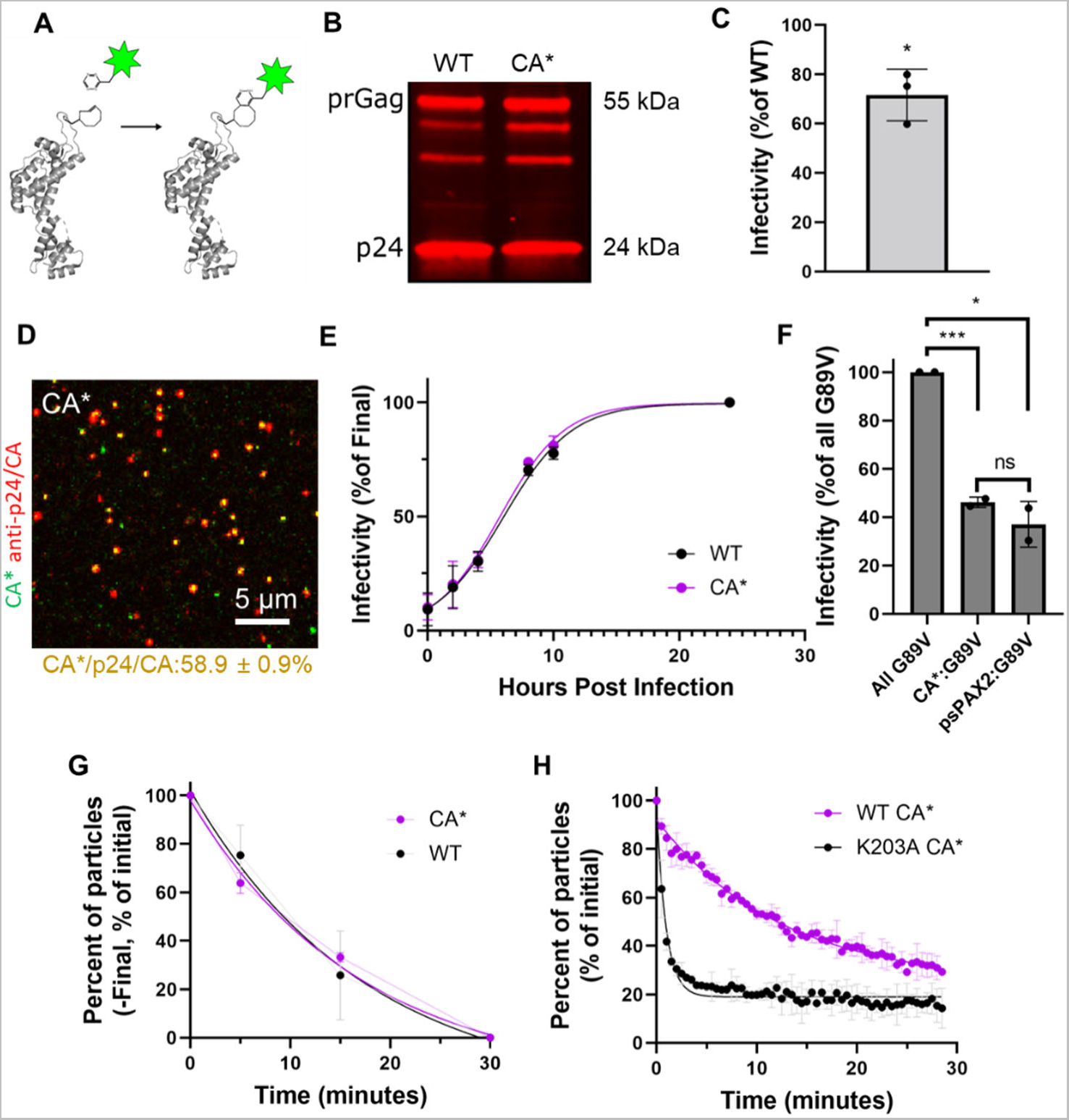
Validation of HIV-1 CA* labeled pseudoviruses. **A)** Schematic illustration of CA protein with the isoleucine-91 site labeled with SiR-tetrazine (green star). **B)** Western blot analysis of labeled viruses. Twenty pg of WT or CA* virus p24 was immunoblotted with human anti-HIV serum antibodies and stained with anti-human LiCor IRDye 800CW for fluorescence detection. Estimated protein molecular weights (kDa) listed next to bands. **C)** Specific infectivity of CA* pseudoviruses measured as firefly luciferase light units normalized to p24 content. CA* specific infectivity is normalized to WT infectivity with mean and standard deviation of 3 viral preparations shown (p = 0.043, unpaired t-test, mean and SD plotted). **D)** Cell-free CA* pseudovirus plated on poly-L-lysine coated coverglass, labeled with 250 nM SiR-tetrazine and immunostained with mouse anti-p24 AG3.0 antibody and anti-mouse AF568 second antibody. Colocalization is calculated between 3 independent preps. Mean and SD are listed. **E)** PF74 time-of-addition nuclear import kinetics assay for WT and CA* pseudoviruses. Here, 2.5 µM PF74 was added to cells at 0, 2, 4, 8, 10, and 24 hpi, (T_50_ for CA* = 6.27 hpi, T_50_ for WT CA = 6.58 hpi, mean and S.D. shown for 3 different viral preps). **F)** TRIMCyp restriction assay to assess the cyclophilin A binding capabilities of CA* pseudovirions. TZM-bl cells expressing TRIMCyp were infected with CA*/WT:G89V pseudoviruses, and the resulting luciferase signal was measured after 48 hours. Firefly luciferase light units were normalized to p24 ELISA content and CA*/WT:G89V ratio was normalized to 100% G89V infectivity (CA*:G89V: p = 0.0008, CA^WT^:G89V: p = 0.01, CA*:CA^WT^: p = 0.32, unpaired t-test, mean and SD shown for 2 different viral preparations). **G)** *In vitro* uncoating assay kinetics curves from CA* and WT pseudoviruses. CA* and WT pseudoviruses were lysed with 100 µg/mL saponin and fixed/immunostained with anti-CA (AG3.0) anti-bodies. Data is from 2 independent preps, mean and S.D. plotted. **H)** Kinetics of *in vitro* uncoating of HIV-1 pseudoviruses containing WT CA* and K203A CA*. The membranes of WT CA* and K203A CA* pseudoviruses were permeabilized with 100 µg/mL saponin. A fraction of particles retaining above-threshold CA* signals are responsible for a non-zero plateau. Data is from two independent viral preps, mean and S.D. are plotted.

To assess the extent of incorporation and labeling of CA* in the context of mixed CA*/WT CA pseudoviruses, virions were adhered to coverslips, labeled with Silicon Rhodamine-tetrazine (SiR-tetrazine), fixed, permeabilized and immunostained for p24/CA. Using this protocol, 58.9% of CA* labeled cores colocalized with p24 signal (Fig. 1D). The less-than-optimal colocalization of CA* with p24/CA may be due to the use of multiple plasmids for transfection, low efficiency of amber codon suppression and/or incomplete click reaction with HIV-1 core-incorporated CA* (see below for additional analyses/validation). Specificity of CA* labeling with SiR-tetrazine is evident from a background level SiR signal after staining of p24 immunostained cores containing WT CA (Fig. S1A). Although recent reports suggest that HIV-1 backbone is click-labeled, unless all amber codons in Vif, Vpu, and Rev are mutated,^59^ the lack of off-target staining of the WT CA vector in our experiments(Fig. 1A) likely reflects the absence of these proteins in budding virions.

We next asked if the intraviral distribution of CA* was unaffected by the mutation and click labeling. Whereas HIV-1 particle contains several thousand Gag polyproteins (>2500),^60, 61^ only about 1500 CA monomers are capsid lattice-associated in mature virions.^60–62^ Using eGFP-tagged CA, it has been reported that permeabilization of the viral envelope results in release of a large portion of labeled CA that was not incorporated into the capsid lattice, trapped within the viral membrane.^25, 32, 44^ In general agreement with these results, exposure of mixed CA*/CA^WT^ pseudoviruses to membrane-permeabilizing saponin released about roughly half of the virus’ CA* signal (Fig. S1B, C).

To test if transport through the nuclear pore of our SiR-tetrazine labeled CA* cores is impaired, we analyzed the nuclear import kinetics by performing a PF74 time-of-addition assay.^26, 41, 63, 64^ At low concentrations, PF74 blocks nuclear import, likely by stabilizing HIV-1 cores, whereas high concentrations of this compound impair reverse transcription and displace imported viral complexes from nuclear speckles.^6, 26, 41, 63, 65^ Thus, the time course of virus escape from a low dose of PF-74 reflects the kinetics of HIV-1 nuclear import.^26^ These experiments revealed that the kinetics of nuclear import of mixed CA*/CA^WT^ pseudoviruses is indistinguishable from that of control pseudoviruses (Fig. 1E).

The cyclophilin A (CypA) binding loop within the capsid lattice interacts with CypA, and likely with Nup358, and plays an important role in HIV-1 infection through modulating capsid interactions with a multitude of host factors (including restriction factors).^5, 8, 66–69^ We therefore asked if the mixed cores containing CA* retain the ability to functionally interact with the owl-monkey TRIMCyp factor, which binds to the CypA binding loop of CA and restricts infection by prematurely degrading the capsid lattice.^19, 21, 44, 70–72^ To indirectly assess TRIMCyp binding, mixed CA* pseudoviruses were produced by co-expressing the untagged G89V CA mutant within the CypA binding loop, which renders the virus resistant to TRIMCyp restriction.^44, 73^ Having G89V CA present in the mixed capsid lattice allows for probing TRIMCyp binding to CA* (or CA^WT^ in control samples), using infectivity readout. CA*/CA^G89V^ and CA^WT^/CA^G89V^ viruses were equally sensitive to TRIMCyp restriction, whereas control viruses containing only pure G89V CA cores were resistant to this factor (Fig. 1F). These results demonstrate that direct labeling of the CypA binding loop at the position I91 does not impair CypA binding to HIV-1 cores containing a mixture of WT CA and CA* and that CA* is incorporated into the capsid lattice, with the CypA binding loop exposed to cytosolic factors.

To ensure that our CA* labeling approach does not impact capsid stability, we compared the stability of CA* and WT cores using an *in vitro* uncoating assay. WT HIV-1 uncoating was measured by immunostaining for p24/CA. Briefly, we lysed coverslip attached pseudoviruses with saponin, fixed the exposed cores at 0, 5, 15, and 30 min after lysis, and immunostained with the anti-HIV-1 p24 AG3.0 antibody which detects mature cores.^74, 75^ There was no significant difference in *in vitro* uncoating kinetics between mixed CA*/WT and unlabeled WT cores (Fig. 1G). This suggests that our CA* labeling protocol does not impact core stability and that, therefore, mixed WT/CA* cores behave like unlabeled WT CA cores. Accordingly, time-resolved imaging of single mixed WT/CA* core uncoating revealed a similar kinetic of loss of CA* labeled cores after saponin lysis (Fig. 1H). Of note, 29% of cores retained post-lysis levels of CA* fluorescence by 30 min, likely representing “closed” capsids^48^ that failed to uncoat under these conditions. In contrast, markedly accelerated uncoating was observed for K203A CA* mutant capsids known to form highly unstable cores that rapidly uncoat after viral fusion or lysis (Fig. 1H and Fig. S1E).^76^ Interestingly, a small fraction of the K203A mutant cores (∼10%) exhibited delayed uncoating over the course of 30 min (Fig. S1F). Also, ∼15% of particles retained very weak but detectable levels of CA* fluorescence by 30 min after saponin lysis (Fig. S1E). Thus, loss of WT/CA* signal at varied times after lysis is primarily associated with the capsid lattice dissociation, whereas residual fluorescence retained by a minor fraction of unstable cores after uncoating may correspond to vRNP-associated CA*.^25, 32, 44^

Since amber codon suppression is not very efficient, the lower mutant Gag polyprotein estimation of the actual CA* and WT CA ratio is unlikely to accurately reflect the respective plasmid ratio. To estimate the actual ratio of CA* and WT CA in virus producing cells, HEK293T/17 cells were transfected with CA* and WT CA viral vectors at a plasmid ratio of 1.5:1 and the resulting protein expression ratio was assessed by Western blotting. To ease the assessment of CA*/WT CA ratio, transfections were carried out in the presence of 200 nM of the HIV-1 protease inhibitor, Saquinavir (SQV).^77^ This blocks Gag cleavage and thereby yields largely a single Gag band, simplifying ratiometric analysis. Analysis of the Gag band intensities demonstrates that there is ∼1.6 times more WT CA produced compared to CA* (Fig. S2). Thus, less than 2:1 ratio of unlabeled to labeled CA rescues HIV-1 infectivity, in sharp contrast to the GFP-tagged CA, which requires 10-15-fold excess of untagged CA.^25, 32^ This result shows that the amber suppression efficiency reduces the TCO-tagged Gag expression in HEK/293T cells relative to untagged Gag by about 2.2-fold.

Together, these results demonstrate that our direct labeling approach minimally affects the virus functionality and the core stability in the context of mixed cores containing WT CA, making it suitable for visualizing the sites and the kinetics of uncoating in target cells.

### HIV-1 core entry into the nucleus is not associated with loss of CA* marker

We took advantage of directly labeled HIV-1 capsid to assess the sites capsid uncoating in cells. GHOST(3) cells^78^ were utilized for HIV-1 nuclear import and uncoating analysis due to their low background staining with SiR-tetrazine compared to HeLa-derived cells and thin nucleus, which facilitates 3D time-lapse imaging. In addition, Tat-driven LTR-GFP reporter expression in these cells allows for detection of infected cells in live cell imaging experiments described below. We made GHOST(3) cells stably expressing SNAP-Lamin B1^26, 44, 79^ (referred to as GHOST-SNAP-lamin cells) to visualize the nuclear membrane by staining with SNAP-reactive dye. Cells were infected with CA* labeled particles pseudotyped by VSV-G that drives HIV-1 pseudovirus entry through endocytic route.^80–82^ Four hours post-infection (hpi), cells were fixed, and the fluorescence intensity distributions of single CA* cores in the cytosol, nuclear membrane, and nucleus were compared.

For meaningful core intensity analysis, it is essential to discriminate between intact virions trapped in endosomes and post-fusion viral cores released to the cytoplasm. This is because fluorescence of free cores is considerably lower than that of intact virions because less than half of viral CA is incorporated into the capsid lattice and is immediately lost upon virus membrane lysis (Fig. S1B) or fusion.^25, 34, 60^ To differentiate intact virions trapped in endosomes from free cytosolic CA* cores, we infected cells in the presence of the fixable membrane marker mCLING-Atto488^83^ that associates with both the viral and cell membranes, as has been demonstrated by imaging pseudoviruses in SupT1 cells^31, 38^ (see also Fig. 2A). Endosome-trapped virions were readily identified by colocalization of CA* fluorescence with mCLING-Atto488 in the cytosol of infected cells (Fig. 2B, yellow circles) and excluded from analysis. Quantification of the intensities of free CA* cores (Fig. 2B, white arrows) in the cytoplasm, at the nuclear membrane, and in the nucleus of fixed cells revealed no significant difference in intensity distributions (Fig. 2C), suggesting there is no detectable loss of capsid protein upon HIV-1 nuclear import.

**Fig 2.**
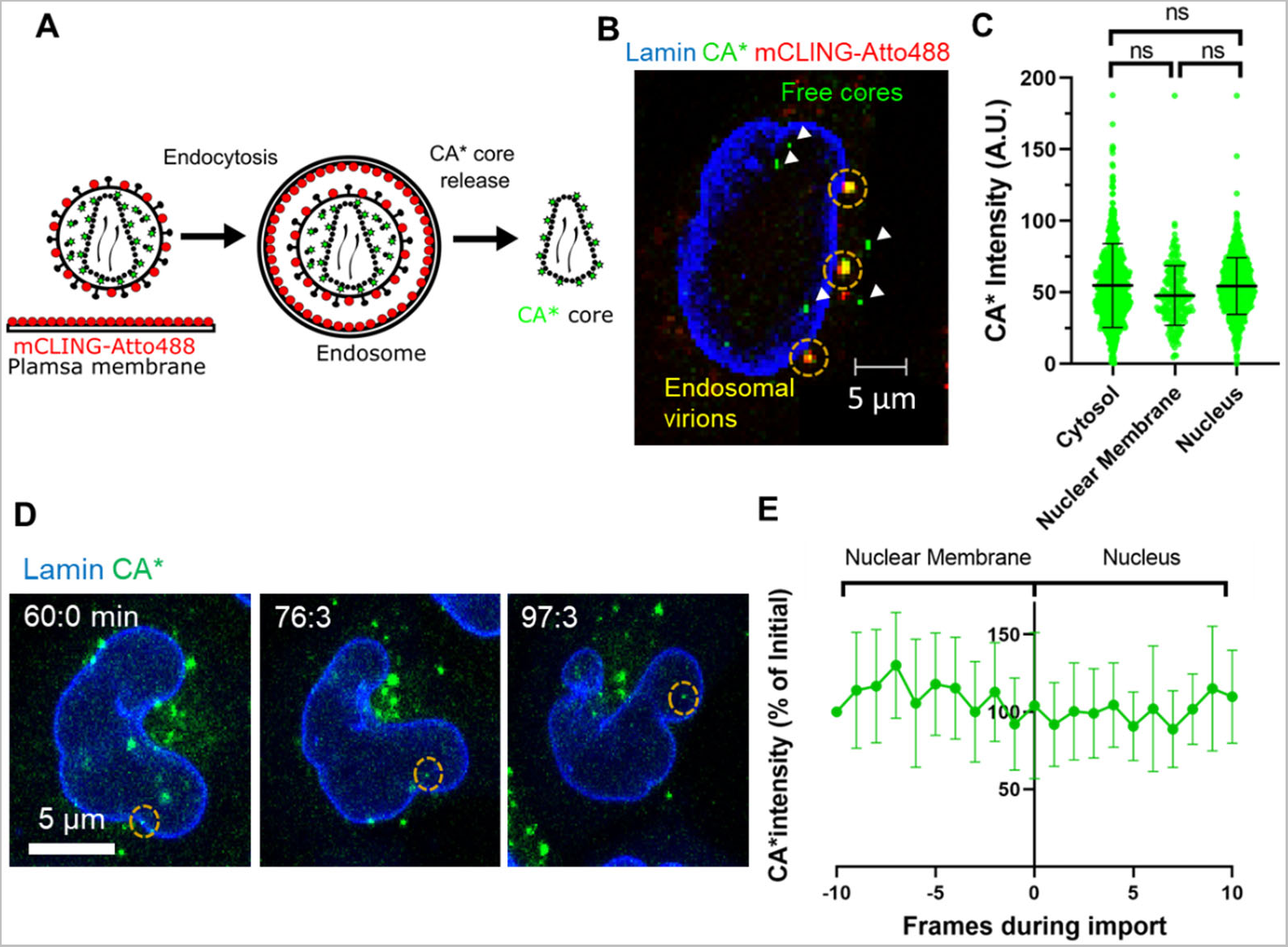
HIV-1 capsid protein is not lost upon nuclear entry into GHOST-SNAP-lamin cells. **A)** Schematic illustration for mCLING-Atto488 labeling of GHOST-SNAP-lamin cells and CA* viral membranes to visualize endosome localized CA* pseudoviruses. **B)** Representative image of infected GHOST-SNAP-lamin cells having endosome localized CA* pseudoviruses (circled in yellow, colocalized with mCLING-Atto488 (red)) and several free CA* cores in the cytosol and nucleus (white arrows). SNAP-Lamin B1 was labeled with SNAP-TMR-Star. Image was taken at 4 hpi. **C)** Fluorescence intensity distributions of cytosolic, nuclear membrane and nucleus localized CA* cores at 4 hpi (Cytosolic: n = 881, Nuclear membrane: n = 299, Nucleus: n = 622) (Cytoplasm:Nucleus: p = 0.995, Nuclear membrane:Nucleus: p = 0.777, Cytoplasm:Nuclear membrane: p = 0.927,Kolmogorov-Smirnov test with optimal binning). **D)** Representative time-lapse micrograph of an individual CA* core entering the nucleus of a GHOST(3) SNAP-Lamin cell at 76.3 minutes post infection, SNAP-Lamin B1 labeled with SNAP-OregonGreen particle of interest is marked by yellow circle. **E)** Ensemble normalized fluorescence intensity profiles of 8 nuclear entry events aligned at the time of nuclear entry.

To extend the analysis of CA* intensity in the cytoplasm and nucleus of fixed cells, we imaged entry of individual CA* cores into the nucleus of living GHOST-SNAP-lamin cells. Importantly, single virus tracking didn’t detect loss of fluorescence intensity upon nuclear import of CA* cores (n=8, Fig. 2D, E, and Movie 1). Although the relatively quick virus docking and nuclear entry in GHOST cells precluded the analysis of CA* intensity prior to and at the time of docking, our fixed cell data support that there is no detectable loss of CA* occurs during passage through the NPC. Collectively, these data suggest that HIV-1 cores with a largely intact capsid lattice may enter the nucleus of GHOST(3) cells.

### HIV-1 core uncoating in the nucleus correlates with productive infection

We sought to visualize the behavior of nuclear CA* cores in live cells and correlate this with productive infection. GHOST-SNAP-lamin cells were labeled with SNAP-reactive dye and infected with CA* viruses at multiplicity of infection (MOI) of 0.2-0.5. Click-labeling of CA* cores were carried out in cells by adding SiR-tetrazine at 0.5 hpi. Single particle tracking in live cells from ∼1.5 to 20 hpi revealed that, after entering the nucleus, HIV-1 cores undergo terminal uncoating manifested in a loss of CA* fluorescence signal (Fig. S3A, B, and Movie 2). On average, CA* signal dropped to a background level at 7.2 hpi (Fig. S3C, n=49), and roughly 84% of the nuclear CA* cores underwent terminal uncoating during our imaging window.

To determine if these nuclear uncoating events correlate with productive infection, Tat-driven GFP-expression in GHOST-SNAP-lamin cells was assessed by infection with CA* GFP reporter pseudoviruses using low MOIs of 0.1. We used GFP reporter pseudovirus in combination with GFP reporter GHOST-SNAP-lamin cells to decrease the waiting time required for detection of GFP expression, since there should be more GFP produced upon Tat expression. We observed nuclear import and subsequent loss of the CA* cores and found that these events correlate with productive infection (Fig. 3A, B, and Movie 3). Approximately 29% of nuclear uncoating events resulted in productive infection. This relatively low percentage of productive events suggests that not all terminal uncoating events culminate in infection but may also be due to the limited imaging window (20 hpi) that may not detect late GFP expression events. While longer live cell imaging is possible, it presents severe practical challenges, such as phototoxicity, photobleaching, and exceedingly large file sizes. The average waiting times for infectious uncoating events was 6.1 hpi (Fig. 3C), while the average time for GFP reporter expression was 12.9 hpi, with an average lag time between CA* uncoating and GFP expression being 6.8 hours (Fig. 3D). As expected from incomplete (∼60%) colocalization of CA* and p24 signals in coverslip-adhered pseudoviruses (Fig. 1B), GFP expression was also observed in cells with no detectable nuclear CA* cores/uncoating events.

**Fig 3.**
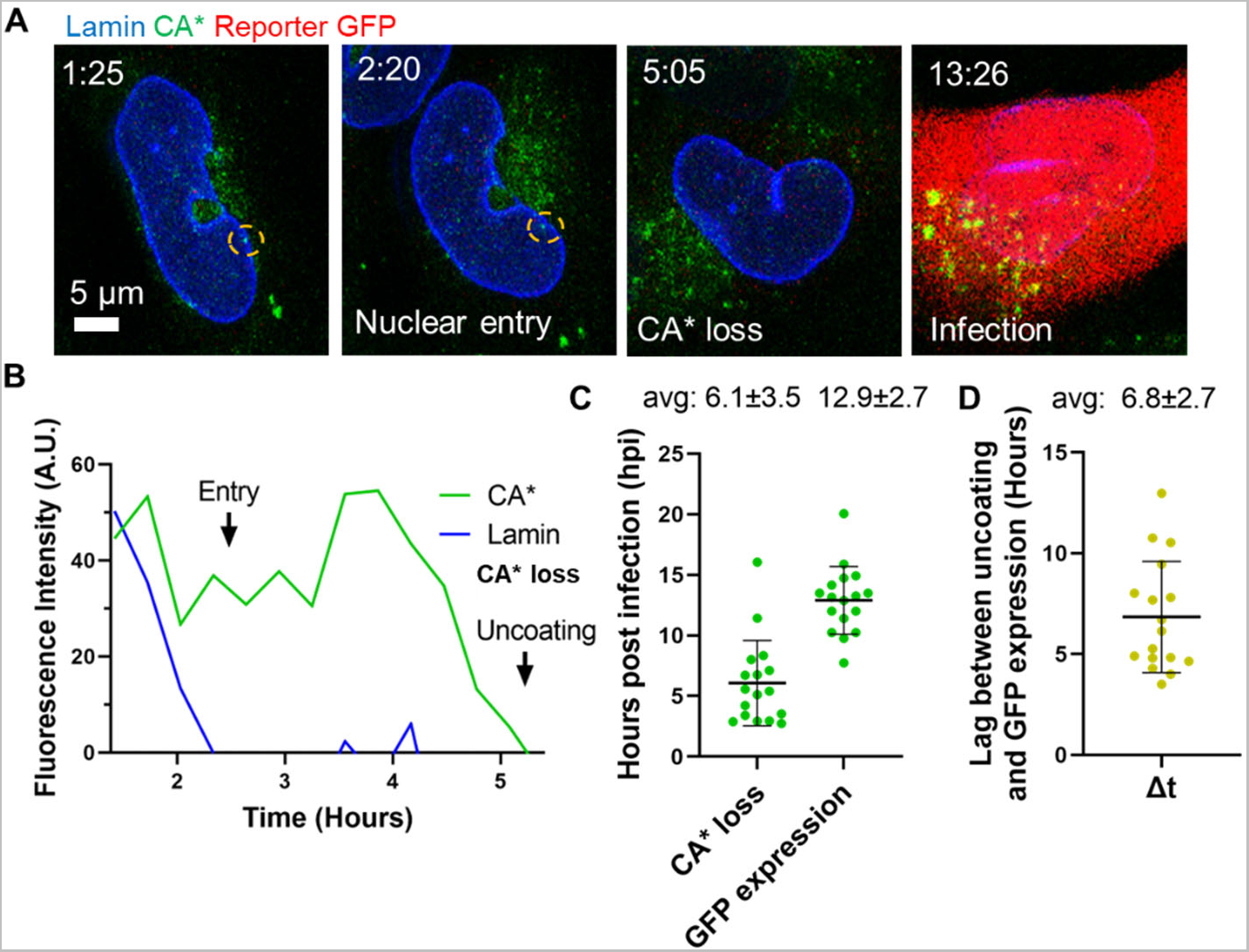
Nuclear CA* cores undergo terminal uncoating that culminates in productive infection. **A)** Representative micrographs of a single CA* core entry into the nucleus, uncoating, and the resulting productive infection (GFP expression). The CA* core is marked by the yellow dashed circle. **B)** Single particle intensity trace of the nuclear CA* core from panel A. Nuclear entry and uncoating are marked with arrows. **C)** Distributions of functional uncoating times and times of detectable GFP reporter expression (n = 17). The average uncoating time and GFP reporter expression time with standard deviations are listed above the graph. **D)** The lag time (Δt) between uncoating and GFP reporter expression in single cells. The average lag time is listed above the graph.

Together, our results suggest that intact HIV-1 cores can enter the nucleus and that nuclear uncoating represents a productive route of infection in cell lines. Note that the difference between the mean uncoating time in the context of reporter GFP expression (Fig. 3C) and for all uncoating events (Fig. S3C) is not significant (Fig. S3D). This modest difference is likely due to insufficient imaging time to track later uncoating events all the way to the reporter GFP expression.

To assess if reverse transcription is completed prior to terminal uncoating GHOST-SNAP-lamin, we measured the kinetics of reverse transcription of CA* cores in GHOST-SNAP-lamin cells through a Nevirapine (NVP) time-of-addition assay to block reverse transcription at different time points.^25, 26^ The time course of escape of CA^WT^ and CA* pseudovirus infections from NVP was very similar, demonstrating that the kinetics of reverse transcription is not affected by CA* labeling (Fig. S4). The shorter T_50_ for virus escape from NVP compared to that for loss of CA* signal in GHOST-SNAP-lamin cells implies that completion of reverse transcription precedes terminal nuclear uncoating leading to productive infection, as suggested previously.^2, 25, 33, 36, 41, 84^.

### HIV-1 uncoating progresses through distinct steps *in vitro*

Our direct CA labeling strategy revealed loss of CA* signal in the nucleus that tends to culminate in productive infection. We next sought to resolve early steps of HIV-1 capsid uncoating preceding the terminal loss of CA*. Single capsid integrity loss has been visualized through incorporation of a fluid phase GFP marker into viral cores.^24, 48, 85^ This approach relies on incorporation of the GagiGFP marker into the viral cores. This Gag construct containing “internal” GFP (iGFP) generates a fluid phase iGFP marker upon virus maturation.^24, 27, 48, 86^ A small fraction of iGFP trapped inside an intact capsid lattice allows for sensitive detection of defect formation in the capsid lattice manifested by release of this marker. *In vitro* uncoating experiments have revealed that a fraction of intact cores, defined as cores that retained residual iGFP signal after viral membrane permeabilization, lost capsid integrity (released residual iGFP) prior to terminal uncoating, measured by loss of cyclophilin A signal, which binds to untagged CA lattice.^48^

To assess both the early defect formation and terminal uncoating of HIV-1 capsid in cells, we incorporated a fluid phase marker into CA* cores. We swapped GFP for YFP in the GagiGFP backbone^47^ to make a Gag-iYFP-Pol construct (hereafter referred to as Gag-iYFP).^24, 27, 47, 48^ iYFP trapped inside the intact HIV-1 cores is released upon the formation of small defects (>4 nm),^87^ thus acting as a core integrity marker (Fig. 4A).^24, 27, 48^ By producing CA*:Gag-iYFP pseudoviruses similarly to the single labeled CA* virions, we achieved 56.2% colocalization of iYFP with CA*, but 91.9% of CA* signal was associated with iYFP (Fig. 4B). Thus, the overwhelming majority of CA* cores contain a fluid phase fluorescent marker. This double labeling approach did not considerably reduce specific infectivity (Fig. 4C). Immunoblotting revealed efficient cleavage of GagiYFP in double-labeled pseudoviruses, as evidenced by a prominent iYFP band (Fig. 4D).

**Fig 4.**
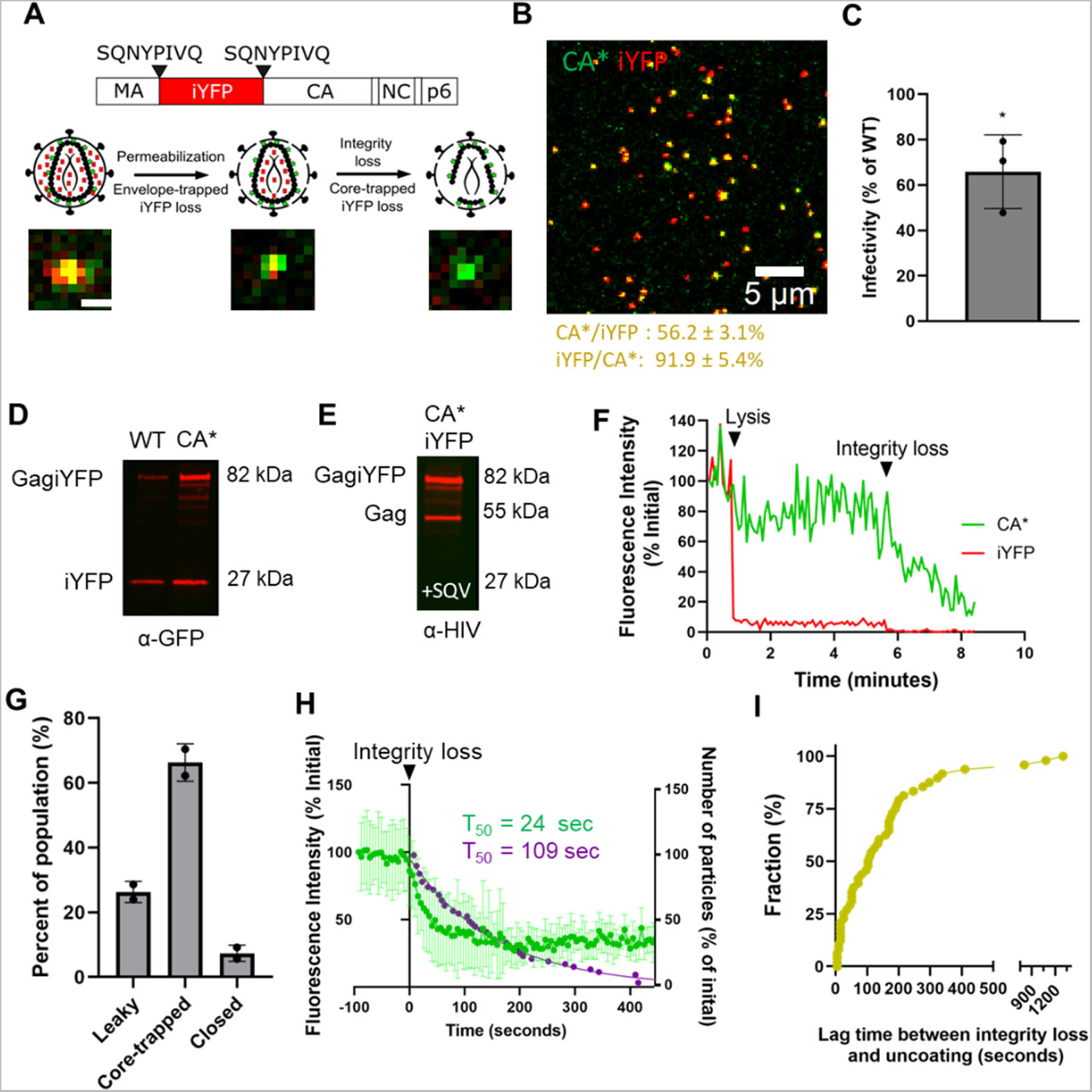
Multi-step uncoating of CA*:Gag-iYFP pseudovirus cores *in vitro*. **A)** Schematic for CA*:Gag-iYFP pseudovirus labeling with the fluid phase marker iYFP for assessing capsid integrity. Schematics and images of cell-free CA*:Gag-iYFP particle integrity loss and uncoating are shown (scale bar is 1 µm). **B)** Representative image of cell-free CA*:Gag-iYFP pseudoviruses labeled with SiR-tetrazine. Percent colocalization is listed below in the yellow font, mean and SD are plotted. **C)** Specific infectivity of CA*:Gag-iYFP pseudoviruses (p = 0.02, unpaired t-test). Mean and SD is shown with 3 different virus preps. Luciferase light units were normalized to CA*:Gag-iYFP p24 content and to WT:iYFP pseudovirus infectivity. **D)** Western blot displaying GagiYFP cleavage in WT:iYFP and CA*:Gag-iYFP lysates (50 pg virus p24 loaded). Anti-GFP living colors antibody was used for iYFP detection. Estimated protein molecular weights (kDa) listed next to bands. We did visualize a somewhat higher uncleaved GagiYFP precursor bands (∼82 kDa) for CA*:Gag-iYFP pseudoviruses, but this is likely due to the reduced efficiency of CA* translation, allowing for more GagiYFP incorporation during virus assembly than WT. There were additional MA-iYFP-CA bands (synonymous with p41 MA-CA band) on the blot, but these are likely due to a small fraction of immature particles present in the population. **E)** Estimation of CA*:Gag-iYFP ratio in HIV-1 cores. Producer cells were treated with 200 nM Saquinavir to block virus maturation, anti-HIV serum antibodies were used for virus detection (50 µg of p24 loaded). Estimated protein molecular weights (kDa) listed next to bands. **F)** Representative intensity trace for two-phase iYFP release phenotype from CA*:Gag-iYFP cores. Arrows and identifying text signify time points of pseudovirus lysis and integrity loss. **G)** Quantification of Gag-iYFP release phenotypes from CA*:Gag-iYFP pseudoviruses in vitro. Each population is normalized to the total particle population. **H)** Ensemble intensity trace plot of coverslip immobilized CA*:Gag-iYFP pseudovirus CA* loss kinetics after iYFP loss. Traces were aligned at iYFP loss (t=0) and normalized to the CA* signal at this time point. Averaged CA* traces were terminated at the time of complete loss of CA* signal. Overlaid on this ensemble averaged curve is the waiting time distribution for terminal loss of CA* after iYFP release (grey symbols). Half-time of CA* loss is shown in green for CA* intensity and half-time for post-iYFP loss uncoating kinetics shown in grey. **I)** The lag times between integrity loss and terminal uncoating from the particles used in Panel H. Kinetics curve is normalized to the final time point.

Similar to the ratio of CA*/CA^WT^ in single-labeled pseudoviruses, transfections of producer cells with a 1.5:1 plasmid ratio (CA*:Gag-iYFP, see Methods) of CA* and yielded the protein ratio of approximately 1:1.6 of CA*/CA^WT^ (Fig. 4E). (Viruses were produced in the presence of 200 nM Saquinavir to block maturation and ease densitometry analysis, as described in Fig. S1C).

We examined disassembly of CA*:Gag-iYFP cores that contained releasable iYFP by lysing HIV- 1 pseudoviruses adhered to coverslips with saponin, as above. Imaging the resulting loss of iYFP and CA* signals revealed three distinct types of iYFP release events: (1) immediate and complete loss of iYFP upon viral membrane permeabilization, likely corresponding to “leaky” cores, (2) a two-phase loss of iYFP: initial release of most of the viral iYFP immediately after addition of saponin, followed by a second loss due to defect formation in intact cores,^24, 27, 48^ and (3) initial iYFP loss upon saponin addition without loss of core-entrapped iYFP, likely representing “closed” cores that failed to uncoat by the end of an imaging experiment.^48^ A representative single particle trace of two-phase release is depicted in Fig. 4F. Examples of leaky and closed traces are shown in Fig. S5A, B. We found that 66.3% of CA*:Gag-iYFP cores exhibit a two-phase iYFP loss, whereas only a minor fraction of cores is “leaky” (26.3%) or “closed” (7.4%) (Fig. 4G). Thus, the majority of HIV-1 cores co-labeled with CA* and iYFP appear to contain intact capsid lattice, in contrast to the Gag-iGFP labeled cores reported to represent a minor fraction of pseudoviral cores.^48^ Interestingly, we often observe gradual loss of CA* content after loss of iYFP, as shown in Fig. 4F (middle panel), suggesting that uncoating *in vitro* may occur through a gradual loss of CA, rather than catastrophic disassembly, which was suggested by an *in vitro* uncoating assay using indirect capsid labeling.^48^

To determine if the fluid phase marker, iYFP, can be released on demand from the CA*:Gag-iYFP labeled cores, coverslip-adhered pseudovirus membrane was permeabilized by saponin in the presence of PF74, which, at higher concentrations, induces core integrity loss but stabilizes the capsid lattice.^25, 27, 41, 48, 65, 88^ At 10 µM, PF74 induces a rapid and complete loss of iYFP from CA* cores, while markedly stabilizing the CA* signal over time (Fig. 4H), in agreement with a previous study.^48^ We can thus utilize PF74 to promote defects in CA*:Gag-iYFP capsids in the cell nucleus in order to ascertain the intactness of these HIV-1 cores.

To investigate whether lattice defect formation leading to iYFP release is associated with detectable loss of CA*, we plotted an ensemble average of 49 single particle intensity traces aligned at the time of the second iYFP release step. Most (∼82%) cores do not exhibit a detectable reduction in the CA* signal at the point of iYFP loss, but CA* intensity gradually decays with a half-time (T_50_) of 23 sec. Interestingly, cores that do not lose all CA* signal after release of iYFP, tend to retain, on average, ∼33% of the initial CA* fluorescence signal (Fig. 4H, green curve). Representative individual intensity traces (Fig. S7) demonstrate a relatively rapid CA* decay after iYFP loss, with only a few CA*:Gag-iYFP cores still retaining CA* by 60 sec after integrity loss. We also determined the kinetics of terminal CA* uncoating after iYFP loss and found the half-time to be ∼109 sec (Fig. 4H, purple curve). The lag time between integrity loss and terminal uncoating for all 49 CA*:Gag-iYFP cores in Fig. 4 is, on average, 150.2 sec (Fig. 4I). These results suggest that, upon defect formation, the capsid gradually loses CA* until a critical threshold (∼33% of the initial CA* core content). From this point, capsids tend to become unstable and undergo terminal loss of the lattice, consistent with the published results.^48, 89^ Importantly, because most cores did not exhibit detectable CA* loss upon loss of capsid integrity, the capsid lattice defect allowing iYFP release *in vitro* is likely to be relatively small.

### Terminal CA* loss of nuclear HIV-1 capsids is preceded by a loss of core-trapped iYFP

Having demonstrated the ability of the CA*:Gag-iYFP virus to report the integrity loss and uncoating events *in vitro*, we investigated if intact capsids (defined as CA* labeled cores containing releasable iYFP) can enter the cell nucleus. To test this, GHOST-SNAP-lamin cells were infected with CA*:Gag-iYFP labeled pseudoviruses and fixed at 4 hpi to determine the amounts of double-labeled and CA* only labeled cores in the nucleus. Over half (60.8%) of the nuclear CA* foci were iYFP positive, suggesting that intact capsids can enter the nucleus of infected cells (Fig. 5A). The nuclear CA* cores lacking iYFP (Fig. 5A’) may, in principle, arise from a loss of integrity event, either before nuclear entry or in the nucleus. Alternatively, these could be “leaky” CA*:Gag-iYFP cores that do not retain iYFP after fusion or lysis (Fig. S5A) or intact cores that were not labeled with iYFP upon production (Fig. 4B). Supporting the latter notion that single-labeled nuclear puncta largely represent cores that failed to incorporate Gag-iYFP upon assembly are the same relative fractions of double-labeled cores in the nucleus and in intact virions adhered to coverslips (after excluding the “leaky” particles from analysis, Fig. 5B). These findings suggest that most nuclear cores in GHOST-SNAP-lamin cells within 4 hpi are intact.

**Fig 5.**
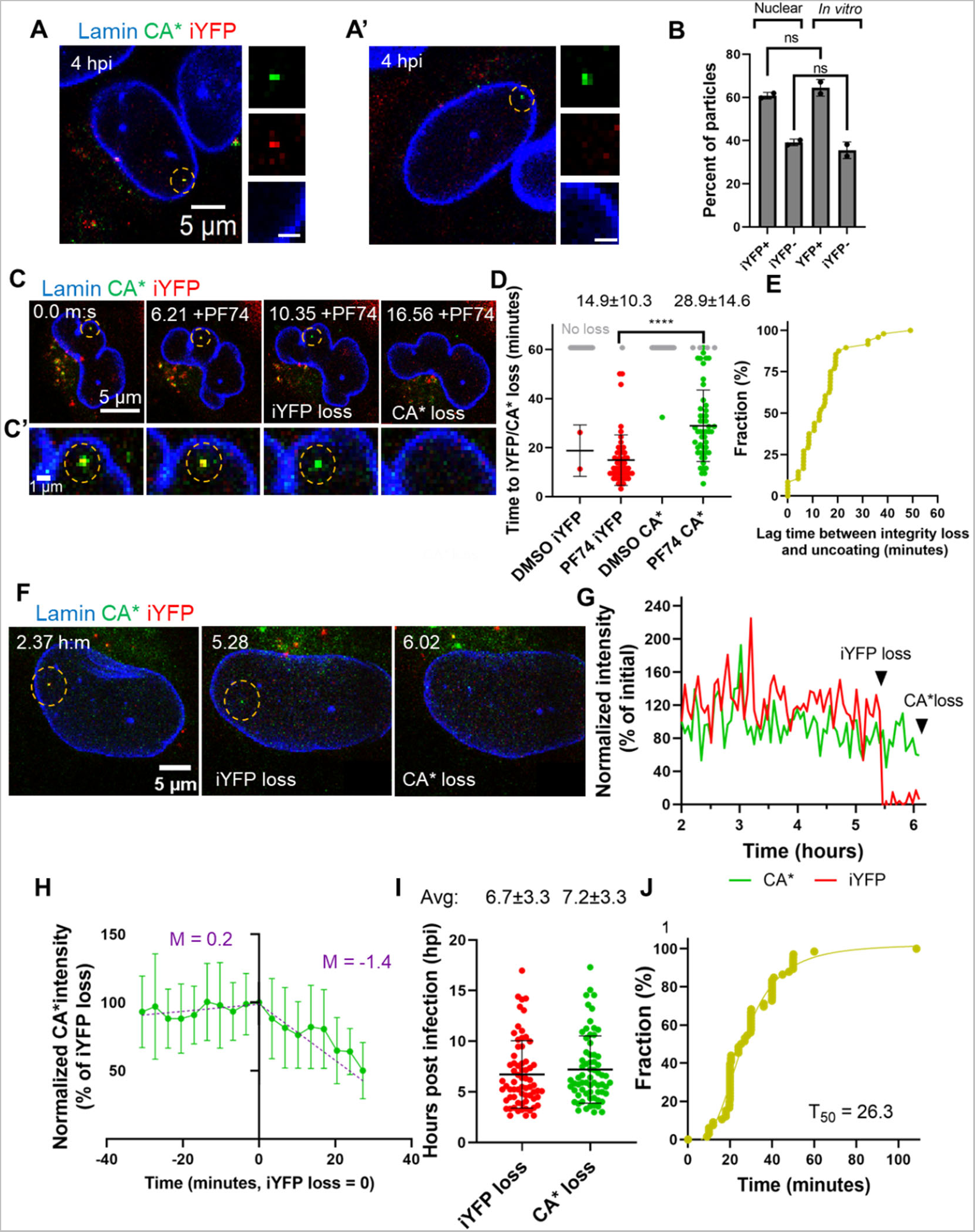
Intact cores enter the nucleus of GHOST-SNAP-lamin cells and undergo consecutive loss of capsid integrity and terminal uncoating. **A) A** representative image showing a nuclear CA*:Gag-iYFP capsid. **A’) A** representative image of a non-intact nuclear capsid lacking iYFP signal. Dashed yellow circles mark the particles of interest. Zoomed in images of the nuclear CA*:Gag-iYFP cores are shown to the right. Scale bar is 1 µm. **B)** Quantification of the fractions of YFP-and YFP+ CA* cores in nuclei and inside saponin lysed viral particles attached to coverslips. Data were normalized to the total number of nuclear particles. Coverslip-attached particles were quantified by subtracting the population of leaky CA*:Gag-iYFP cores (see Figure 4D, G, and H). Means and SD are plotted. **C)** Live cell micrograph of a nuclear CA*:Gag-iYFP core in GHOST-SNAP-lamin cell treated with 10 µM PF74. The particle of interest is marked with the yellow dashed circle. **C’)** Zoomed in section of the live cell micrograph from panel C. **D)** The kinetics of iYFP and CA* loss in PF74 and DMSO treated GHOST-SNAP-lamin cells (p <0.0001, Mann-Whitney rank sum test). Grey dots represent CA*:Gag-iYFP particles that remain after 1 hour of imaging. Mean and SD are plotted. Average time and SD are listed above each PF74 column. **E)** Cumulative distribution plot for the lag time between integrity loss and PF74-induced capsid degradation in the nucleus of cells. **F)** Representative live cell micrograph of CA*:Gag-iYFP integrity loss and uncoating in the nucleus of an infected GHOST-SNAP-lamin cell. Particle of interest is highlighted by the yellow dashed circle. Integrity loss and uncoating are marked by white text within the image. **G)** Representative single particle intensity traces for the CA*:Gag-iYFP core from panel F. Integrity loss and uncoating are marked by arrows. **H)** Ensemble intensity trace plot demonstrating CA* uncoating dynamics after iYFP loss. Blue text lists the slope for the per-integrity loss and post integrity loss ensemble average for the CA* intensity trace. Traces longer than 30 minutes were cut off at 30 minutes to improve curve fitting. **I)** Waiting times of all CA*:Gag-iYFP integrity loss and uncoating events within the nucleus. Mean and SD are plotted. Average integrity loss and uncoating times are displayed above each column with standard deviations. **J)** Cumulative distributions of the lag time between integrity loss and uncoating of single particles. Mean and S.D. are plotted.

To ensure that the nuclear cores contain a releasable pool of iYFP, we induced the release of iYFP by treating infected GHOST-SNAP-lamin cells with PF74, similar to *in vitro* experiments (Fig. S6). Addition of 10 µM PF74 to cells at 3 hpi, induced a loss of iYFP from CA* cores, while the CA* signal was retained for an additional several minutes (Fig. 5C, and Movie 4). The majority (88.5%) of iYFP-positive CA* cores released iYFP upon PF74 addition, while retaining CA* signal. Only 1.6% of CA*:Gag-iYFP cores were not sensitive to PF74 treatment, resulting in no observable loss of iYFP or CA* during the 1-hour incubation with PF74. In contrast, there were only two integrity loss events and one CA* loss event that was visualized during this imaging window in DMSO treated cells, showing that PF74 potently triggers integrity loss in nuclear HIV- 1 cores. We did detect several instances (9.8%) of iYFP and CA* being lost within the same image frame, but this is likely due to our limited temporal resolution (∼2 minutes) that is insufficient to resolve short lags between the two events. By tracking 50 nuclear double-labeled cores, we find that the average time for iYFP loss after PF74 treatment is 14.9 minutes, followed by CA* loss after a significant lag of 13.9 minutes, on average (Fig. 5D, E). Interestingly, nuclear CA* cores become much more mobile upon PF74 treatment, consistent with our previous work, which has reported increased core mobility of nuclear cores displaced from nuclear speckles and subsequent degradation after PF74 treatment.^6^ Thus, delayed loss of CA* signal after release of iYFP upon PF74 addition can be attributed to degradation of HIV-1 cores displaced from nuclear speckles.^6^

We next asked if release of iYFP is temporally linked to terminal loss of CA* in the nucleus. To visualize capsid integrity loss and uncoating, we tracked 67 CA*:Gag-iYFP labeled pseudoviruses in live GHOST-SNAP-lamin cells between ∼1.5 hpi and ∼22 hpi. Representative single particle images and fluorescence traces show a loss of capsid integrity (iYFP release) followed by terminal uncoating (loss of CA*, Fig. 5F, G, and Movie 5). Importantly, for all visualized CA*:Gag-iYFP labeled nuclear cores, loss of core integrity culminated in terminal uncoating. The average time for capsid integrity loss was 6.7 hpi, whereas subsequent terminal uncoating occurred at 7.2 hpi. The latter time is very close to that of terminal uncoating of single-labeled CA* cores (compare Figs. S3C and 5I), implying that double-labeling of cores does not affect their nuclear import or stability. The average lag time between iYFP release and terminal loss of CA* in GHOST-SNAP-lamin cells is 29 minutes (Fig. 5J), suggesting that loss of integrity of the capsid lattice may be an intermediate step of terminal uncoating. Interestingly, no detectable loss of CA* is evident after averaging the CA* intensity traces (n=18) aligned at the time of iYFP loss, indicating that the initial defect responsible for iYFP release is not associated with a significant loss of CA from the lattice. While no instant loss of CA* at the time of iYFP release is detected, the CA* signal gradually decays from that point on with a slope of 1.4 %/min (Fig. 5H). A gradual loss of CA* following an initial lattice defect formation is consistent with our *in vitro* uncoating results (Fig. 4I), except that gradual uncoating in the nucleus is a much slower process compared to *in vitro* uncoating (T_50_ = 24 sec).

We next asked if reverse transcription influences the lag time between integrity loss and uncoating within the nucleus. We visualized the kinetics and efficiency of integrity loss (iYFP release) and uncoating (CA* loss) in GHOST-SNAP-lamin cells in the presence of the reverse transcriptase inhibitor NVP (10 µM). The integrase strand-transfer inhibitor, Raltegravir (RAL, 10 µM), was used as a control since integration occurs after capsid disassembly. Inhibition of reverse transcription delayed capsid integrity loss and uncoating by roughly 2.5 hours each. The average time for integrity loss and uncoating in the presence of DMSO is 6.2 and 6.8 hpi, as compared to 8.8 and 9.3 hpi, respectively, in NVP treated cells (Fig. 6A). NVP treatment also causes a 19.1% decrease in the probability of CA* loss observed within our imaging window (Fig. 6B), suggesting that reverse transcription plays a role in the uncoating process, in agreement with.^2, 19, 25, 33, 36, 39, 41, 65, 90, 91^ The equally delayed iYFP release and loss of CA* in the presence of NVP imply that reverse transcription promotes initial defects in the capsid lattice, without significantly affecting terminal uncoating. Importantly, the lag time between integrity loss and uncoating was not affected by DMSO, NVP, or RAL treatment (Fig. 6C). This indicates that reverse transcription plays a role in initiating capsid integrity loss but does not have a major role in the terminal uncoating process after integrity loss.

**Fig. 6.**
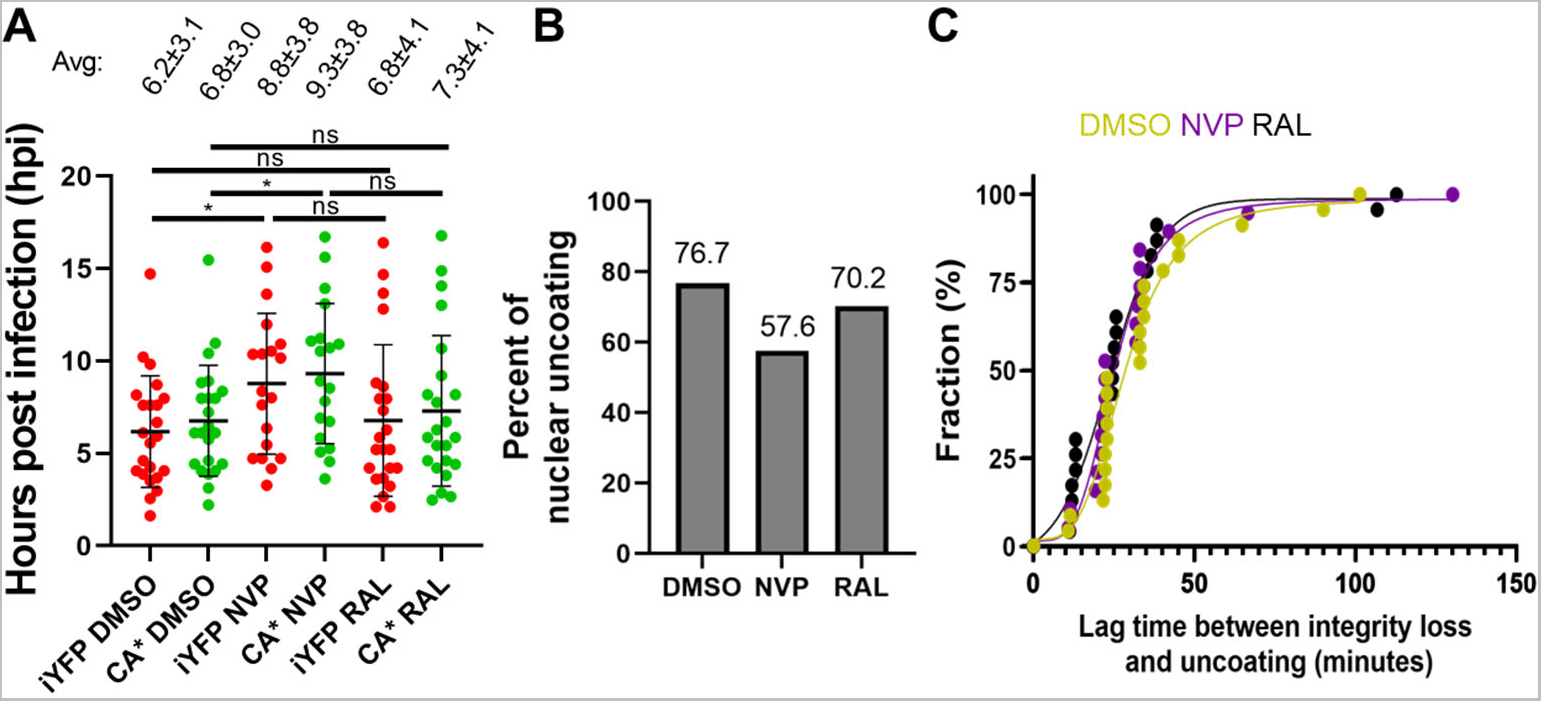
Reverse transcription accelerates capsid integrity loss but does not affect the lag time between integrity loss and uncoating. **A)** Waiting times for all CA*:Gag-iYFP integrity loss and uncoating events with DMSO, 10 µM NVP, and 10 µM RAL treatment. Mean and SD are plotted. Average integrity loss and uncoating times are displayed above each column with standard deviations. **B)** Uncoating efficiency of DMSO and NVP treated cells, with 19.1% reduction in NVP treated CA*iYFP cores, with minimal effect with RAL present. **C)** Cumulative distributions of the integrity loss and terminal uncoating lag times in DMSO, NVP, and RAL treated cells.

### HIV-1 undergoes two-step uncoating in different cell types

To generalize the observed HIV-1 uncoating dynamics in the nucleus of transformed GHOST cells, we examined the core integrity loss and CA* disassembly in differentiated THP-1 macrophages-like cells, which model the nuclear environment of physiologically relevant cells. The HIV-1 core integrity loss and uncoating events in the nucleus of THP-1 derived cells were difficult to visualize within our imaging window due to the SAMHD1 activity in these cells, which reduces the dNTP pool^26, 92, 93^ and slows down reverse transcription.^93, 94^ We were nonetheless able to detect single core uncoating events in these cells by overnight imaging, but, due to the slow rate of reverse transcription, the visualization of uncoating was likely limited to early events. Like in GHOST cells, HIV-1 cores in the nucleus of THP-1 macrophages underwent two-phase uncoating – release of iYFP followed by terminal uncoating (loss of CA*) (n=10, Fig. 7A, and Movie 6). The representative single particle intensity traces (Fig. 7B) depict the temporal relationship between loss of capsid integrity and terminal uncoating occurring 26.4 min after release of iYFP. For particles that do uncoat during our imaging time, the average times of integrity loss is 11.3 hpi, and the subsequent terminal uncoating events are at 11.8 hpi (Fig. 7C). This is consistent to the functional uncoating start time in THP-1 macrophages deduced by blocking the HIV-1 import through the nuclear pore at different times post-infection and exposing nuclear cores to a high concentrations of PF74.^36^ These results demonstrate that sequential core integrity loss and uncoating occur in the nucleus of physiologically relevant cells. Importantly, the average observed lag time between capsid integrity loss and terminal core uncoating was 31 minutes (Fig. 7D), which is strikingly close to that in GHOST-SNAP-lamin cells (Fig. 5H). Thus, the lag time for integrity loss and loss of CA* appears independent of the cell type, suggesting that terminal uncoating after an initial defect formation may be intrinsically regulated by the capsid lattice and its inter-hexamer/pentamer interactions.^95^

**Fig 7.**
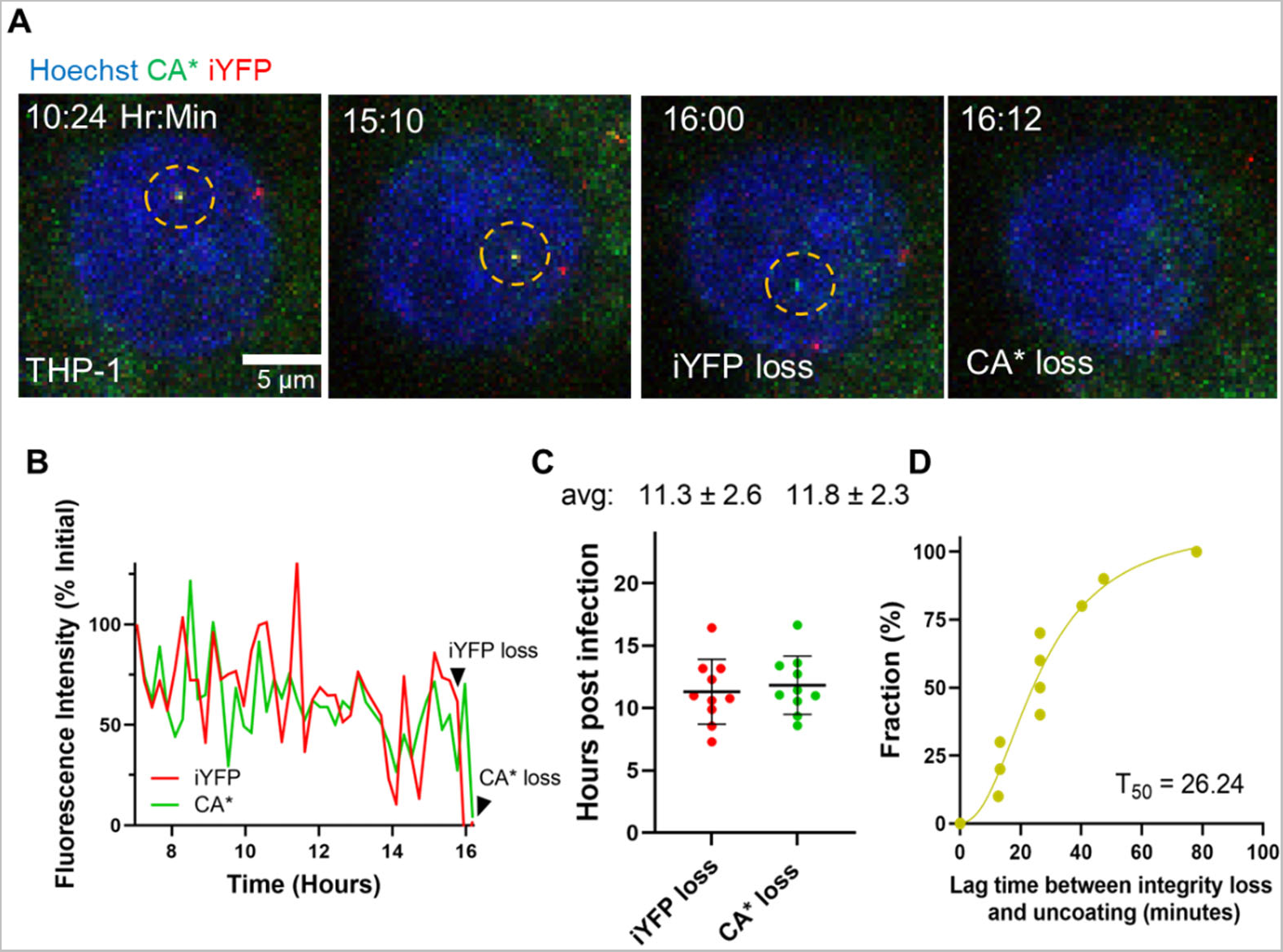
Loss of capsid integrity precedes terminal uncoating in THP-1 derived macrophages. **A)** Representative live cell imaging micrograph of THP-1 derived macrophage cell (stained with 2 µM Hoechst33342) containing an single CA*:Gag-iYFP core undergoing loss of integrity (iYFP) and subsequent terminal uncoating (loss of CA*). Integrity loss and uncoating listed in frame and CA*:Gag-iYFP core is highlighted by yellow dashed circle. **B)** Single particle intensity trace of the CA*:Gag-iYFP core from panel A. Integrity loss and uncoating times are marked with arrows. **C)** Integrity loss and uncoating times in THP-1 derived macrophages. Mean and SD are plotted. Average values and standard deviations are listed above each column. **D)** Cumulative distribution plot for Δt values for loss of integrity and uncoating for single particles.

## Discussion

Here, we implemented a minimally invasive dual labeling scheme to visualize the loss of integrity and uncoating of single HIV-1 capsid in live cells, in the context of productive infection. Virions containing a ∼1.6:1 mixture of WT and amber mutant CA exhibit nearly unaltered infectivity, maturation, capsid stability, host-factor binding, and nuclear import. This direct labeling strategy, combined with incorporation of a fluid phase marker of capsid integrity (iYFP), reveals the dynamics of single HIV-1 uncoating in the nucleus of living cells. By comparison, nuclear import of cores solely composed of CA click labeled at the position A14, using the amber codon suppression approach, is strongly delayed.^34^ Nonetheless, the fixed cell-based fluorescence intensity analysis of single A14-labeled cores in the cytoplasm and the nucleus showed no significant difference in CA* signals. This result supports the notion that intact or nearly intact HIV-1 cores enter the nucleus, in full agreement with our data (Fig. 2A-C). Through live cell imaging, we consistently observe HIV-1 uncoating proceeding through at least two distinct steps – loss of iYFP, apparently through a local defect in the capsid lattice, followed by disassembly of capsid lattice (terminal uncoating) after a significant delay.

Our laboratory has previously developed the tetrameric CypA-DsRed (CDR) marker that incorporates into viral particles by tightly binding to HIV-1 CA at the cyclophilin A binding loop.^26, 30, 49^ We have observed a terminal loss of CDR at the nuclear pore roughly 20 minutes prior to nuclear entry.^26, 30, 49^ Thus, loss of core-bound CDR prior to nuclear import appears distinct from loss of iYFP or CA* in the nucleus. A possible explanation is that CDR is displaced from the cyclophilin A binding loop by Nup358, which has been reported to bind to this loop,^5, 8^ or other CA-interacting nucleoporins during transport through the nuclear pore. Although the inner diameter of the nuclear pore is ∼60 nm,^38^ it is filled with a dense meshwork of FG nucleoporins that likely pose a significant barrier for HIV-1 core import,^13^ leaving little room for additional proteins coating the capsid. This dense meshwork may also play a role in the nuclear import delay of the amber codon mutant at the A14 position due to the high levels of dye conjugation to pure A14 cores.^34^

The size of an initial defect in the capsid lattice, and, therefore, the extent of CA loss upon iYFP release from the core is not known. In principle, even loss of a single hexamer (roughly 9-10 nm)^56, 57^ is sufficient to release iYFP (∼4 nm). The lack of detectable decrease in CA* signal at the time of iYFP release (Fig. 4I, 5H) is consistent with a relatively small defect in capsid lattice. However, given the rather low signal/background ratio (∼3:1) in live cell imaging settings, a loss of as much as 25% of CA* may remain undetected. We thus cannot rule out the possibility that larger defects may be responsible for iYFP release and that the vDNA is released through these defects. However, the fact that iYFP release and subsequent degradation of nuclear HIV-1 cores induced by PF74 are associated with inhibition of productive infection (Fig. 1E & 5D)^6, 27, 63, 88^ argues against this possibility. Although it is unclear whether PF74-induced defects are of the same size as local defects forming naturally prior to uncoating, the inhibition of infection by PF74 is inconsistent with the vDNA release from HIV-1 cores prior to terminal uncoating, which would allow vDNA integration and productive infection.

Several lines of evidence support the ability of the HIV-1 capsid lattice to tolerate relatively large defects without fully disassembling.^65, 90, 91, 95, 96^ Atomic force microscopy experiments suggest that HIV-1 uncoating *in vitro* is driven by reverse transcription and proceeds through a localized rupture at regions of high curvature of the capsid lattice.^65, 90, 91, 95, 96^ This conclusion is supported by course-grained and all-atom molecular dynamics simulations revealing that capsid integrity loss/breakage during reverse transcription occurs in regions of high curvature.^95^ Our results support the notion that reverse transcription accelerates integrity loss and thereby increases the efficiency of uncoating, without significantly reducing the lag between integrity loss of uncoating (Fig. 6A, B, C). However, inhibition of reverse transcription lowered the probability of uncoating without stopping this process.^25^ This suggests that there may be a yet unknown host factor within the nuclear speckle/nucleoplasm that controls capsid stability.

Recent CLEM data revealed the presence of broken HIV-1 capsids in the nucleus that appear to lack the vRNP-associated density but maintain the hexagonal CA lattice.^33, 38^ Zila and co-authors observed release of vDNA from nuclear HIV-1 INmScarlet labeled fluorescent cores and correlated these foci with broken/intact cores within the nucleus using CLEM and electron tomography. These results suggest that the HIV-1 capsid does not fully disassemble in the nucleus, but rather undergoes breakage/cracking and retains a major portion of the lattice. Based on the visualization of damages capsids, it has also been proposed that the HIV-1 capsid lattice undergoes remodeling during nuclear import.^42–44^ However, retention of the releasable iYFP marker after nuclear import is inconsistent with extensive capsid remodeling, unless this process occurs without any defect formation. Furthermore, our observation that the CA* signal is lost upon terminal uncoating ∼30 minutes after integrity loss is inconsistent with the capsid breaking model^33, 38^ and supports the full uncoating model.

It has been shown that low levels of CA protein can associate with the vRNP complex *in vitro* and *in vivo*.^44^ Single particle tracking and biochemical evidence supports the association of untagged CA with the vRNP components (NC, IN, and RT). Our control experiments revealed that only a small fraction (∼15%) of unstable K203A CA* cores retained detectable low-level fluorescence after uncoating *in vitro* (Fig. S1D and SE), which may correspond to vRNP associated CA*. This very low residual CA* signal is unlikely to be detected in the nucleus, since a typical signal/background ratio in our live cell experiments is ∼3:1. Our intensity analysis with HIV-1 cores demonstrates that cores in the cytosol, nuclear membrane, and nucleus have very similar CA* intensity distributions (Fig 2A-C), suggesting that the CA* signal observed in the nucleus represents intact or nearly intact cores and not the residual vRNP associated CA* pool. We therefore interpret loss of CA* signal in the nucleus as terminal uncoating of HIV-1 capsid and not loss of the vRNP associated CA pool.

Our results agree with the nuclear uncoating models proposed previously,^25, 27^ which posits that intact HIV-1 capsids enter the nucleus, and that subsequent uncoating occurs through terminal disassembly leading to productive infection (detected based upon visualizing the sites of viral gene transcription). However, the experiments leading to this uncoating model did not employ double-labeled pseudoviruses that enabled us to dissect the dynamics of single HIV-1 uncoating in the nucleus. Instead, two separate series of experiments have been performed using eGFP-CA labeled cores or cores single-labeled with a fluid phase marker.^25, 27^ When using a fluid phase marker for core integrity, loss of the capsid lattice has been visualized indirectly, using fluorescently tagged CPSF6 that accumulates around intact nuclear cores.^27^ This indirect detection of capsid uncoating after loss of a fluid phase marker suggested that uncoating occurs 1-3 minutes after integrity loss.^27^ In contrast, our dual labeling approach that enables direct visualization of these two uncoating steps reveals a rather long (∼30 min) lag between loss of capsid integrity and terminal uncoating.

The marked delay in capsid uncoating after the initial loss of capsid integrity supports the need for an additional trigger to disassemble the capsid and release the pre-integration complex. Notably, the invariance of this lag time in osteosarcoma and macrophage cell lines suggests the involvement of virus-intrinsic processes or conserved cellular processes occurring after the initial loss of capsid integrity to trigger terminal capsid disassembly.

The notion of multi-step HIV-1 uncoating in the nucleus is consistent with our *in vitro* uncoating data (Fig. 4F, I, J). Our results are in full agreement with the report that single HIV-1 uncoating *in vitro* progresses through "capsid opening” (release of a fluid phase marker), followed by gradual loss of CA protein and culminating in catastrophic loss of capsid visualized based on an indirect staining with fluorescent cyclophilin A.^48^ Unlike HIV-1 uncoating in cells, the half-life of capsids *in vitro* after virus membrane permeabilization is under 12 min, and the lag time between integrity loss and uncoating is only around 70-100 sec. Although this lag time has been reported to nearly double in the presence of cell lysate or IP6^48, 89^. Uncoating of post-fusion HIV-1 cores in cells takes hours, and the lag time from initial core opening to capsid disassembly is much longer (∼30 min, Fig. 5F-J) than in cell free experiments (∼2.5 min, Fig. 4H). Such marked difference in the core stability likely reflects HIV-1 interactions with host factors in the context of intact cells.^5, 35, 97^

The emerging model is that the intact HIV-1 capsid is essential for host factor interactions that facilitate nuclear import and transport to nuclear speckles, protection of the vRNA/vDNA from host-immune surveillance within the cytosol and nucleus, and retention of requisite viral enzymes for completion of reverse transcription and integration into host genome. Premature loss of CA protein in the cytosol/nuclear membrane may expose the viral genome to innate cellular responses, including cGAS.^98^. Our study provides critical insights into the dynamic process of capsid uncoating within the nucleus of cells. We hypothesize that intact capsids enter the nucleus, and uncoat *via* a multi-step mechanism that progresses through: (1) the formation of a small defect, (2) gradual CA loss till critical threshold is reached, and (3) terminal disassembly (Fig. 8). Our results suggest that the HIV-1 capsid can tolerate significant loss CA, but, once the loss of CA reaches a critical point, catastrophic capsid disassembly occurs. These results demonstrate that HIV-1 capsid stability is tightly regulated and that small defects can trigger a catastrophic loss of the lattice, albeit after as significant delay. Future studies will aim to delineate the nuclear site(s) and intermediates of uncoating resulting in the release of the pre-integration complex.

**Fig 8.**
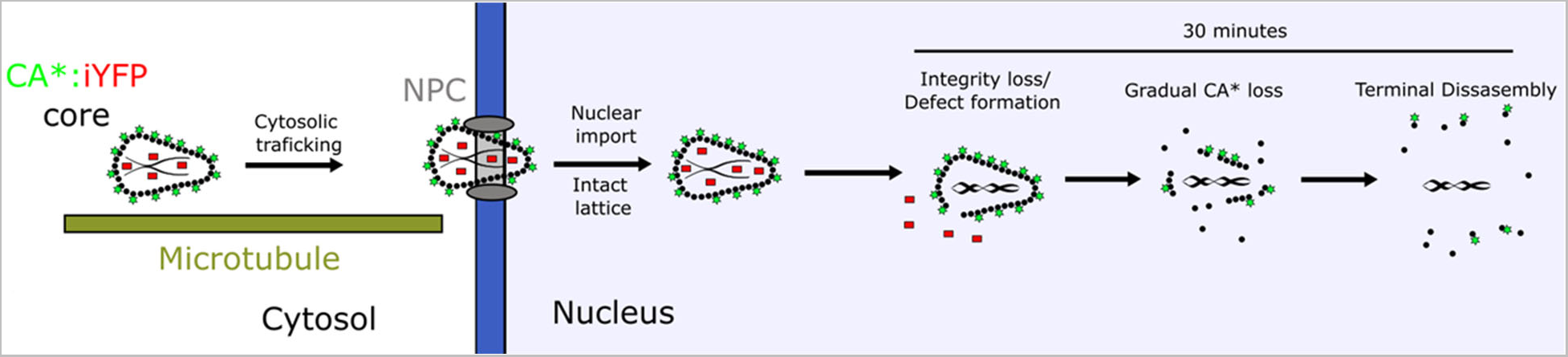
A model for HIV-1 capsid uncoating that progresses through small defect formation, gradual CA* loss, critical lattice fraction preceding terminal disassembly of the lattice in the nucleus.

### Limitations of Study

The limitation of our current labeling technique lies in imperfect colocalization of CA* and p24/CA signals in the mixed pseudovirus cores. As a result, we cannot rule out the possibility that unlabeled cores uncoat outside the nucleus or that these uncoating events do not proceed through at least 2 distinct steps. However, given the minimal effects of our direct CA labeling strategy on virus’ infectivity, interactions with host factors, and core stability, we believe that the observed uncoating of double-labeled viruses reflects productive uncoating of unlabeled HIV-1 cores. Additional limitations include a relatively low signal-to-background ratio and limited temporal resolution that preclude the detection of a small fractional CA loss and/or of any residual CA signal after terminal uncoating. Further optimization of direct CA labeling and imaging techniques should help overcome the above shortcomings.

## Conclusions

Imaging of single HIV-1 cores containing a mixture of WT CA and directly labeled CA (CA*) and co-labeled with a fluid phase marker (iYFP) provides a wealth of information regarding the cellular sites, dynamics, and extent of capsid uncoating. The observation that both loss of iYFP and, after a significant delay, loss of CA* occur in the nucleus supports the conclusion that HIV- 1 uncoats in this cellular compartment and that uncoating is a multi-step process. The lack of detectable loss of CA* at the time of iYFP release, followed by gradual decay of the CA* signal, suggests that uncoating progresses through a distinct intermediate step of small defect in the capsid lattice that triggers a cascade of events leading to terminal uncoating. The invariance of the lag time between loss of iYFP and CA* observed in divergent cell types supports the notion that processes triggered by the initial loss of capsid integrity are intrinsic to the viral core or determined by core interactions with ubiquitous nuclear factors.

## Materials and Methods

### Plasmids

The CA* amber TAG stop codon mutation was introduced into the pNL4-3 R-E-eGFP and pR9ΔEnv plasmid by amplification of the MA-CA region between the BssHII and SpeI restriction sites (forward primer: 5’ – TGCAGCaagcttGCGCGCACGGCAAGAGG – 3’, reverse primer: 5’ – GAGTCGggatccACTAGTAGTTCCTGCTATGTCACTTCCCCTTGG – 3’) and ligating the domain into the pUC19 cloning vector (Addgene plasmid # 50005), using the T4 ligase ligation kit (New England Biolabs, Cat# M020S0). Quikchange mutagenesis primers were constructed to mutate the isoleucine-91 site to an amber stop codon (TAG, forward primer for mutation: 5’-cagtgcatgcagggccttaggcaccaggccagatg-3’, reverse primer for mutation: 5’- catctggcctggtgcctaaggccctgcatgcactg-3’). The mutated partial MA-CA domain was re-inserted into the BssHII and SpeI sites of pNL4-3 R-E-eGFP or pR9ΔEnv to construct the pNL4-3ΔEnv R-E-eGFP or pR9ΔEnv CA* plasmids, respectively. WT backbone vectors were derived using the pNL4-3 R-E-eGFP construct, as described in.^6, 26, 30^ The pR9ΔEnv K203A CA* plasmid was constructed by digesting the pR9ΔEnv CA* plasmid with the BssHI and SpeI restriction enzymes and ligating this MA-CA fragment into the pR9ΔEnv K203A plasmid (A kind gift from Dr. Christopher Aiken). G89V CA was derived from the pR9ΔEnv G89V HIV-1 backbone vector (a kind gift from Dr. Christopher Aiken).^99^ The Gag-iYFP pNL4-3ΔEnv was derived from the Gag-iGFP vector^47^, developed by Naoyuki Kondo (Kansai Medical). The pMD2.G VSV-G vector encoding for VSV-G envelope glycoprotein was obtained from John Naughton at the Salk Institute. The peRF1 E55D and PylRS NES AF amber codon suppression plasmids were a kind gift from Dr. Edward Lemke (Johannes Gutenberg University).^55^

### Cell Lines and Reagents

HEK293T/17 cells (ATCC, Manassas, VA), HeLa-derived TZM-bl cells ectopically expressing CD4 and CCR5^100^ (NIH AIDS Reference and Reagent Program, ARP-8129, contributed by J Kappes, X Wu), TZM-bl cells expressing TRIMCyp^101^ (kind gift from Dr. Jeremy Luban, University of Massachusetts), GHOST(3) CXCR4/CCR5 cells (NIH AIDS Reference and Reagent Program, contributed by V. KewalRamani & D. Littman) were grown in a complete high-glucose Dulbecco’s Modified Eagle Medium (DMEM, Mediatech, Manassas, VA) supplemented with 10% heat-inactivated Fetal Bovine Serum (FBS, Sigma, St. Louis, MO) and 100 U/mL penicillin-streptomycin (Gemini Bio-Products, Sacramento, CA). THP-1 monocytic cells (ATCC, Manassas, VA) were grown in complete Roswell Park Memorial Institute growth media (RMPI, Gibco) supplemented with 10% heat inactivated FBS and 100 U/mL penicillin/streptomycin. Growth medium for HEK293T/17 cells was additionally supplemented with 0.5 µg/mL Neomycin (Mediatech).

GHOST-SNAP-lamin cells were obtained by transduction with VSV-G pseudotyped viruses encoding for SNAP-Lamin B1 transgene. Transduction efficiency was assessed by visual inspection of SNAP-Lamin expression using the SNAP-Cell® TMR-Star (NEB, #S1905S). SNAP-TMR-Star staining was performed as manufacturer’s protocol.

The non-canonical amino acid transcyclooctene lysine (TCO*A, Cat# SC-8008) was purchased from SiChem (Bremen, Germany) and dissolved at 100 mM in 15% DMSO, 0.2 M NaOH, and stored at -20 °C. The clickable Silicon Rhodamine-Tetrazine (SiR-tetrazine, Cat#SC008) dye was purchased from Spirochrome (Thurgau, Switzerland), dissolved in methanol at 1 mM, aliquoted in 20 µL evaporating with argon gas, and stored at -20 °C. Aliquots of SiR-tetrazine were redissolved in 20 µL of DMSO at a final concentration of 1 mM prior to use. The mCLING-Atto488 (Cat# 710 006AT3) membrane marker was purchased from Synaptic Systems (Gottingen, Germany), dissolved in de-ionized water at 50 nmol/mL and stored at -80 °C. The Bright-Glo^TM^ luciferase assay kit was purchased from Promega (Madison, WI). Primary IgG antibodies specific to Lamin B1 (#ab16048) and the secondary IgG donkey anti-rabbit AF405 conjugated antibody (#ab175651) were purchased from Abcam (San Francisco, A). The SNAP-Cell® TMR-Star and Oregon Green® were purchased from New England Biolabs (NEB, #S1905S, #S9104S). Hoechst33442 was purchased from Thermo Fisher (Cat# 62249). HEPES 1M solution (Cat# SH30237.01) was purchased from Cytiva Life Sciences (Marlborough, MA). Phosphate buffered saline containing Mg^2+^/Ca^2+^ (dPBS+/+) and Mg/Ca-free (dPBS-/-) were purchased from Corning (Mediatech, Manassas, VA). The mouse anti-GFP A.V. living colors antibody was purchase from Clontech (Cat# 632381). The Alexa Fluor 568 goat anti-mouse IgG H+L antibody was purchased from ThermoFisher (Cat# A11004). The capsid binding inhibitor PF74 was purchased from Sigma-Aldrich (CAT# SML0835). The HIV-1 protease inhibitor Saquinavir (ARP-4658) and the integrase inhibitor Raltegravir (HRP-11680) were obtained through the NIH HIV Reagent program, (contributed by DAIDS/NIAID). The mouse IgG anti-p24/CA specific antibody AG3.0 (donated by Dr. J. Alan),^102^ reverse transcriptase inhibitor Nevirapine (ARP-4666), integrase inhibitor raltegravir (HRP-11680), protease inhibitor (ARP-4658), Anti-HIV-1 antibody specific to p24/CA hybridoma183 (ARP#1513) and Human HIV-IgG serum (ARP#3957) were all received from the NIH HIV reagent program.

### Pseudovirus Production and Characterization

To produce CA*:Gag-iYFP dual-labeled viruses, the pNL4-3ΔEnv R-E-eGFP CA* or pR9ΔEnv I91* backbone (1.5 µg), pNL4-3ΔEnv Gag-iYFP (1.0 µg), pMD2.G VSV-G (0.2 µg), PylRS NES AF (1.0 µg), and peRF1 E55D (1.0 µg) were transfected into single wells (3.5 cm) of the Corning Costar 6-well cell culture plate containing HEK293T/17 cells using the JetPrime Transfection reagent (VWR, Randor, PA) according to the manufacturer’s protocol. Virions labeled only with CA* were produced using pNL4-3ΔEnv R-E-eGFP CA*/pR9ΔEnv I91* backbone (1.5 µg), pNL4-3 R-E-eGFP CA^WT^/ pR9ΔEnv CA^WT^/psPAX2 (1.0 µg) and the pMD2.G VSV-G, PylRS NES AF, and peRF1 E55D vectors at the same ratio as for the CA*:Gag-iYFP pseudovirus production. For TRIMCyp restriction assays, control pseudoviruses were produced using pR9ΔEnv G89V (2.0 µg) with the pMD2.G VSV-G, PylRS NES AF, and peRF1 E55D vectors at the same ratio as for the CA*:Gag-iYFP pseudovirus production. K203A CA* pseudoviruses were produced using pR9ΔEnv K203A CA* (1.5 µg), and pR9ΔEnv K203A (1.0 µg) with the pMD2.G VSV-G, PylRS NES AF, and peRF1 E55D vectors at the same ratio as described. Complete DMEM transfection medium was supplemented with 250 µM TCO*A, pre-diluted 1:5 in 200 mM HEPES. To produce eGFP-CA labeled pseudoviruses, the pHGFP-WT-BglSL-GFP (1.8 µg), pHGFP-eGFP-CA-BglSL-GFP (0.2 µg), and pMD.2 VSV-G (0.2 µg) were transfected into HEK293T/17 cells as described above.

For production of transducing viruses for SNAP-Lamin B1, the pLENTI-SNAP-Lamin B1 HIV-1 transfer vector (2 µg), psPAX2 (Addgene #12260, Kind gift from Didier Trono, 1 µg), and pMD2.G VSV-G (0.2 µg) was transfected into HEK293T/17 cells using the JetPrime transfection kit as per manufacturer’s recommendation.

After 16 hours of transfection, the growth medium was replaced with 2 mL of complete DMEM without phenol red, supplemented with 250 µM TCO*A. After the media change, the cells were incubated for an additional 32 hours at 37 °C in 5% CO_2_. After a total of 48 hours, the pseudovirus-containing supernatant was collected, filtered through a 0.45 µm filter, concentrated 10x using the Lenti-X lentivirus concentrator (Clontech Laboratories, Inc. Mountainview, CA), resuspended in dPBS+/+, and stored at -80 °C. Pseudovirus p24 content was quantified from viral preparations using the p24 ELISA assay.^103^ The multiplicities of infection (MOIs) were determined in GHOST-SNAP-lamin cells by examining the percent of GFP positive cells after 48 hours post-infection with VSV-G CA* pseudoviruses using a BioTek Cytation 3 imaging plate reader (instrument use is courtesy of Dr. Baek Kim).

### Pseudovirus SPIEDAC Click-Labeling

TCO*A labeled CA*:Gag-iYFP and CA* pseudovirus cores were labeled in cells post/during virus-cell fusion. Viral cores were labeled at 30 min and 60 min post-infection for GHOST-SNAP-lamin and THP-1 cells, respectively, by incubating cells with 250 nM SiR-tetrazine in Fluorobrite DMEM (Gibco, CAT#A1896701) without FBS for 20 minutes. Cells were washed once with dPBS+/+, and further incubated in complete Fluorobrite medium.

### Single-Round Infectivity and Restriction Assay

Virus infectivity was measured by luciferase reporter activity in TZM-bl cells or TZM-bl cells expressing exogenous TRIMCyp. TZM-bl cells were plated in glass-bottom 96-well plates and infected 12-20 hours after plating with VSV-G pseudotyped viruses at 1:100, 1:1,000, & 1:10,000 dilutions. Virus-cell binding was facilitated by centrifuging cells with virus-containing solutions for 30 minutes at 1550xg and 4 °C. At 48 hpi, cells were lysed, and the luciferase activity was quantified using the Bright-Glo^TM^ luciferase substrate (Promega) using the supplier’s protocol. Raw luciferase values were normalized to pseudoviral p24/CA content to derive specific infectivity. GHOST-SNAP-lamin cells were plated and infected with VSV-G pseudotyped HIV-1 CA* and CA^WT^ virions. Pseudoviruses were centrifuged onto cells by the same method described in the luciferase protocol. After 30 minutes post-infection, pseudovirus containing cells were labeled with SiR-tetrazine as described above and incubated in complete DMEM for an additional 47 hours. At 48 hpi, cells were stained with 2 µM Hoechst33442 for 10 minutes, washed with dPBS, then fixed with 4% PFA (Electron Microscopy Sciences, Cat# 1570-S) in dPBS for 20 minutes. Nine adjacent fields of view were acquired for each well, and the number of GFP positive cells and cell nuclei, stained with Hoechst33442, were quantified using the cell segmentation protocol within the BioTek plate reader software.

### Western Blotting

HIV-1 CA* and CA^WT^ pseudoviruses suspension (20 pg p24) was heated to 95 °C, reduced with ß-mercaptoethanol, then loaded ono a 4-15% precast SDS-PAGE gel, with the Precision Plus Protein^TM^ kaleidoscope ladder (BioRad, Cat# 1610375). The contents of the SDS-PAGE gel were transferred to a 0.45 NC nitrocellulose membrane (Cytiva, Cat# 10600012) and blocked with 10% skimmed milk in PBST (PBS+/+ with 0.1% tween-20). The membrane was then incubated with the HIV-IgG human serum antibodies (1:2,000) and/or the mouse monoclonal anti-GFP living colors IgG antibody (1:1,000) for one hour at room temperature or overnight at 4 °C. After three 5-minute washes with PBST, the membrane was incubated with the secondary IgG donkey anti-mouse IRDye® 800CW conjugated antibody (1:10,000) (Li-Cor, Cat# 926-32212) or goat anti-human IRDye® 800CW (1:10,000) (Li-Cor, Cat# 926-32232). The membrane washes were repeated, and the membrane was maintained in de-ionized water. The membrane was visualized using a Li-Cor Oddessy Clx fluorescent gel imager, using the 700, and 800 emission detectors at a resolution of 169 µm for band visualization.

For CA*:WT and CA*:Gag-iYFP stoichiometry determination, 50 µg of lysates of HEK293T/17 cells transfected with 1.2 µg CA* or 0.8 µg CA^WT^/iYFP backbone (with proper amber suppression machinery) in the presence of 200 nM Saquinavir were immunoblotted, as described above. Protein concentrations in cell lysates were quantified using the Micro BCA^TM^ protein assay kit (ThermoFisher, Cat# 23235). On a separate nitrocellulose membrane, GAPDH was immunoblotted with the Rabbit anti-GAPDH polyclonal antibody (1:1000, ThermoFisher, Cat# PA1-987), and visualized with the IRDye 800CW Donkey anti-Rabbit IgG secondary antibody (Licor, Cat# 9926-32213).

### THP-1 Macrophage Differentiation

For visualization of CA*:Gag-iYFP uncoating in THP-1 derived macrophage cells, 100,000 THP- 1 cells were treated with 125 nM phorbol 12-mysristate 13-acetate (PMA, Sigma-Aldrich, CAT# P1585) in complete RPMI-1640 medium for 24 hours. After 24 hours, PMA containing medium was removed, and cells were incubated in complete RMPI-1640 medium for an additional 24 hours.

### Single Virus Imaging *In Vitro*

eGFP-CA, Unlabeled CA* and CA^WT^ pseudoviruses were centrifuged onto poly-L-lysine treated #1.5 8-well chambered cover glass slides (ThermoScientific, Cat# 155409) for 15 minutes at 1550xg, 4 °C. The chambers were washed 1x with dPBS+/+, blocked with 20% FBS in dPBS+/+ for 30 minutes at room temperature. After blocking, 250 nM SiR-tet was added to the immobilized pseudoviruses in 10% FBS in dPBS+/+ for 20 minutes at 37 °C, followed by an additional washing step with DPBS+/+. The fluorescently labeled pseudoviruses were imaged on a Zeiss LSM880 confocal microscope using a C-Apo 40x/1.3 NA or 63x/1.4 NA oil objective or on a GE Healthcare DeltaVision widefield epifluorescence microscope. Multiple fields of view were acquired using the highly attenuated 488, 561, 633 nm laser lines, with respective EYFP, TMR, and Alexa Fluor 647 emissions bands collected using the Gallium-Arsenide (GaAsP) spectral detector for the LSM880. With the DeltaVision system, the respective FITC and Cy5 excitation and emission windows were used and detected with an EMCCD camera. For *in vitro* uncoating assays, viral membranes were permeabilized with 100 100 µg/mL saponin (Riedel de-Hähn) after 5-10 image frames acquired every 5 seconds or after 1 frame being imaged every 30 seconds, and viruses were imaged for 25 min in the presence of permeabilizing agents.

### Immunofluorescence and Fixed Cell Imaging

For fixed-cell imaging, GHOST-SNAP-lamin cells were plated onto collagen or poly-L-lysine treated #1.5 8-well chamber slides and infected with CA* or CA^WT^ at MOI of 0.2-0.5 *via* centrifugation. At 30 minutes post infection, cells were labeled with 250 nM SiR-tetrazine in Fluorobrite DMEM without FBS then washed with dPBS+/+, as described. GHOST-SNAP-lamin cells were labeled with SNAP-TMR-Star or SNAP-Oregon Green® for 20 minutes before infection, using the supplier’s protocol. Cells were incubated at varying times post infection and fixed with 4% PFA for 20 minutes at room temperature. PFA was quenched with 20 mM Tris in dPBS+/+, and cells were either imaged immediately in dPBS+/+ or subjected to immunofluorescence labeling. For the latter protocol, cells were permeabilized with 0.25% Triton X-100 (Sigma-Aldrich, Cat# X100-100ml) for 5 minutes at room temperature, washed three times with PBST, and blocked with 20% FBS in PBST. After blocking, rabbit anti-LaminB1 antibodies (1:1,000, Abcam #ab16048) in blocking buffer were added to the cells for 1 hour at room temperature, washed three times with PBST, followed by the addition of the anti-rabbit AF405 secondary antibody (1:1,000, Abcam #ab175651) for 1 hour at room temperature. After secondary antibody incubation, the cells were washed with PBST three times and imaged in dPBS+/+.

For experiments using mCLING-Atto488, GHOST-SNAP-lamin cells were labeled with 2 µM mCLING-Atto488 for 5 minutes prior centrifuging with pseudoviruses in the presence or mCLING-Atto488, as described above. Virus-decorated cells were washed with DMEM to remove unbound viruses and excess mCLING-Atto488 and incubated in DMEM until labeling with SiR-tetrazine, as described above.

### Live-cell Imaging of HIV-1 Infection

Single HIV-1 capsid integrity loss, uncoating, and subsequent infection were visualized by live cell imaging using confocal microscopy. 50,000 GHOST-SNAP-lamin cells were plated onto collagen coated 35 mm #1.5 cover glass imaging dishes (Matek, CAT# P35G-1.5-10-C) and infected at varying MOIs (0.1-0.15 for correlating uncoating with infection, 0.2-0.5 for visualization of uncoating for a shorter time, without correlating with infection) with VSV-G pseudotyped CA* NL4-3 eGFP particles co-packaged with either pNL4-3 CA^WT^ eGFP/psPAX2 or NL4-3 GagiYFP. Virus binding to cells was enhanced *via* centrifugation (1550xg, 30 minutes, 4 °C). Prior to infection, GHOST-SNAP-lamin cells were labeled with SNAP-Cell® TMR-Star or Oregon Green®, as per supplier’s protocol. After centrifugation, cells were labeled with SiR-tetrazine, as described. Cells were then washed with dPBS+/+ and incubated in Fluorobrite DMEM with 10% FBS and 20 mM HEPES in a Zeiss LSM880 microscope equipped with temperature-, humidity-and CO_2_-controlled environmental chamber. Cells were maintained in 10 µM aphidicolin (Sigma-Aldrich, CAT# A0781-5mg) to block cell division. 3D time-lapse imaging was performed with a Zeiss LSM880 laser scanning confocal microscope using a C-Apo 40x/1.3NA oil immersion objective. All live cell imaging experiments were conducted using the highly attenuated 488, 514, 561, and 633 nm laser lines with the Gallium-Arsenide (GaAsP) spectral detectors with 0.21 or 0.23 µm pixel size and the pinhole size of ∼4.0 Airy Units (AU).

To track nuclear import of CA* cores, cells were imaged at 1 hour post-infection, for a total of 2- 3 hours every ∼1-2 minutes using a tile scan of 16 adjacent fields-of-view with 5-7 Z-stacks spaced by 0.7 µm. To track relationship between uncoating and productive infection, live cell imaging was performed from 1.5-2 hpi up to 24 hours post-infection, with 100 adjacent fields-of-view acquired using 9-11 Z-stacks spaced by 0.5 µm every ∼18 min. Axial drift was compensated using the Carl Zeiss DefiniteFocus module. To track integrity loss and uncoating, 49 adjacent fields-of-view were imaged every 5-10 minutes for a total of 20 hours post-infection with 9-11 Z-stacks spaced by 0.5 µm.

To track the integrity loss and uncoating events in THP-1 derived macrophages, 100,000 THP-1 cells were differentiated with 125 nM PMA on Poly-L-Lysine, as described above. 2 µM Hoechst33342 stained THP-1 macrophages were infected with 200 pg CA*:Gag-iYFP pseudoviruses and labeled with SiR-tetrazine, as described above. Live cell imaging experiments began at ∼1.5 or ∼7 hpi. Sixty-four adjacent fields-of-view were acquired every ∼12 minutes.

### Image analysis

To analyze co-labeling efficiency and *in vitro* uncoating kinetics of CA*, iYFP labeled and p24- immunotsained HIV-1 cores, three random fields of view were processed with the ComDet c.0.5.5 ImageJ plugin. Single pseudovirus particles were detected with 3-pixel size spot detection and 3- 5 intensity thresholding. The *in vitro* uncoating kinetics was plotted as the number of CA* and iYFP labeled particles over time normalize to the initial number of double-positive particles. Immature pseudoviruses and stable cores that did lose integrity or undergo uncoating were excluded from analysis.

To analyze mCLING-Atto488 labeled GHOST-SNAP-lamin cells for CA* fluorescence intensity distributions, the ICY bioanalysis (http://icy.bioimageanalysis.org) protocol development tool was used. Each individual channel (CA*, Lamin, mCLING-Atto488) was used for segmentation. mCLING-Atto488 colocalized CA* cores were eliminated with the subtract ROI tool by segmenting mCLING-Atto488 foci and removing them from the image. Nuclear membrane-localized cores were detected by HK-means detection in ICY with a binary mask overlayed onto the Lamin signal and used as the detection mask for the spot detector plugin. Intranuclear cores were detected by convexifying Lamin ROIs and calculating the space within the Lamin fluorescent signal using the erode ROI processor. Cytosolic cores were detected by subtracting nuclear and nuclear membrane fluorescence from the images and using the spot detector plugin tool to detect all non-nuclear and mCLING-Atto488(-) cores. All CA* ROIs had an additional 2 pixels dilated from their detection foci to quantify and subtract the local background fluorescence intensity for each particle.

Single particle tracking and live cell imaging experiments were analyzed using the Zeiss Zen Black software to visually determine the time of intranuclear uncoating and integrity loss events. Live cell micrographs were annotated to list the uncoating/integrity loss times with the text editor tool within the Zen Black software. To track the single particle fluorescence intensity over time, the ICY spot tracking/manual tracking plugin was used due to the high mobility and nuclear rotation of GHOST-SNAP-lamin cells. Micrographs were first reduced to 2D by cropping cells of interest, extracting Z-planes containing particles of interest, and using maximum intensity projections for single particle tracking using ICY. The ICY track manager intensity profile and background subtraction processer was used to make intensity traces. Intensity traces were normalized to the initial fluorescence intensity value, when applicable. Ensemble intensity trace analysis was conducted by normalizing fluorescent CA* values to the time of iYFP loss *in vitro* and within the nucleus of cells.

GHOST-SNAP-lamin cells have high degrees of mobility. To correct this motion in live cell imaging experiments, the TrackMate V7 ImageJ tool was used. Cells of interest were imported into the TrackMateV7 tool, and individual Hoechst33342 and Lamin fluorescent nuclei were detected using the LoG detector with 15-30 µm object segmentation. Individual segmented nuclei were tracked using the simple LAP tracker tool with varying gap-closing parameters for each cell. Individual cell tracks were selected, and the track stack processing tool was used to superimpose each time frame to center the cell-of-interest in the middle of the movie’s field-of-view. The subsequent movies were then processed through ICY or Zeiss Zen Black software.

### Statistical analysis

Statistical analysis was carried out using GraphPad Prism (9.3.1) or MatLab (R2021a). All infectivity-based data were analyzed using the Student’s unpaired t-test of Prism. Fluorescence intensity data with non-normal distributions were analyzed with Prism using the Mann-Whitney rank sum non-parametric test. For the mCLING-Atto488 intensity distribution experiments, custom MatLab scripts were used to analyze data with a non-parametric Kolmogorov-Smirnov test with optimal binning to account for large differences in population sizes. Uncoating curves were quantified in Prism using the non-linear curve fitting processor with the one-phase decay equation (*Y* = (*Y*0 – *Plateau*) * exp(-*K* * *X*) + *Plateau*. PF74 time-of-addition experiments were curve-fitted using the non-linear regression curve fitting processor with the Sigmoidal equation (*Y* = *Bottom* + (*X*^*Hillslope*) * (*Top - Bottom*)/(*X^HillSlope^* + *EC*50*^HillSlope^*). Linear slope fitting for ensemble intensity analysis was performed with the *Y* = *mx* + *b*. Statistical significance was assigned as p < 0.05 (*), p < 0.01 (**), p < 0.001 (***), p < 0.0001 (****) respectively. T_50_ values for nuclear uncoating lag time kinetics were curve fitted with the sigmoidal equation Y = 100 * (X^HillSlope)/(EC50^HillSlope + (X^HillSlope)).

## Acknowledgments

We gratefully acknowledge Naoyuki Kondo (Kansai Medical), Christopher Aiken (Vandervilt University), Vinay Pathak (NCI, Fredrick), Jeremy Luban (University of Massachusetts Medical School), and Edward Lemke (Johannes Gutenberg University) for their gift of reagents. We are grateful to Mariana Marin and Gokul Raghunath (Emory University), Ashwanth Francis (Florida State University), and René Gifford (Vanderbilt University) for helpful suggestions and critical reading of the manuscript. We also thank members of the Melikyan lab, Hui Wu, Matthew Prellberg, Manav Kumar, for technical assistance, and Yen-Cheng Chen, You Zhang, and Gokul Raghunath for advice on microscopy applications. This work was supported by the NIH R01 AI129862 and R01 AI148382 grants to GBM and by the Behavior of HIV in Viral Environments (B-HIVE) Center U54 AI170855 grants.

## Supporting Information Available online

This manuscript contains supporting information consisting of 7 supplemental figures (Figs. S1-S7) that provide informative content supporting the main figures and text and 1 supplemental table (Table S1). We also include 6 supplemental movies (Movies S1-S6) to demonstrate the dynamics of HIV-1 nuclear entry and uncoating in live cells.

---

**Fig. S1.**
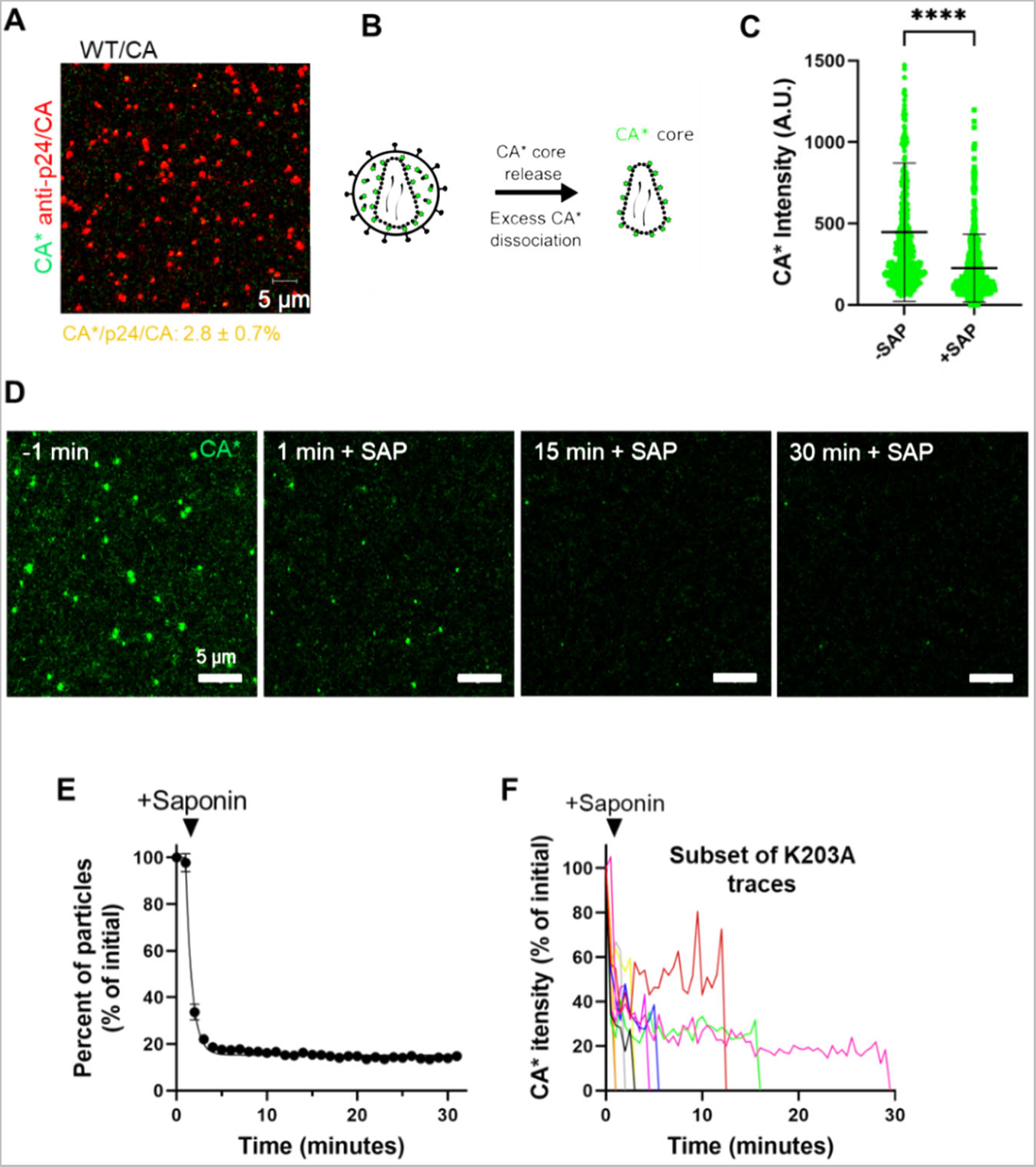
Additional validation of CA* pseudoviruses. **A)** Cell-free WT pseudovirus plated on poly-L-lysine coated coverglass, labeled with 250 nM SiR-tetrazine and immunostained with mouse anti-p24 AG3.0 antibody and anti-mouse AF568 second antibody. Mean and SD are listed. **B)** Representative illustration demonstrating the loss of non-lattice incorporated CA* upon fusion/lysis. **C)** Fluorescence intensity of cell-free viruses labeled with SiR-tetrazine and permeabilized with 100 µg/mL saponin (p <0.0001, Mann-Whitney rank sum test). **D)** Representative micrographs of K203A CA* core uncoating *in vitro* using stringent imaging conditions (0.09 x 0.09 µm pixel size). Saponin (100 µg/mL) was added to coverslip adhered K203A CA* pseudoviruses, and virus membrane permeabilization and loss of CA* monitored by single particle tracking. Particles retaining CA* fluorescence at 1 min after saponin addition (1 min +SAP) represent K203A cores that did not uncoat immediately upon lysis. **E)** *In vitro* K203A CA* pseudovirus uncoating kinetics imaged as in panel D. Mean and SD from 2 independent experiments are plotted. Incomplete loss of CA* signal from a small fraction of labeled pseudoviruses by the end of experiment (30 min) is responsible for the non-zero plateau in the uncoating kinetics. **F)** A small subset (∼15%) of single K203A CA* particle intensity traces from the *in vitro* uncoating experiments in Fig. 1H, demonstrating a delayed loss of K203A CA* signal after saponin lysis.

**Fig. S2.**
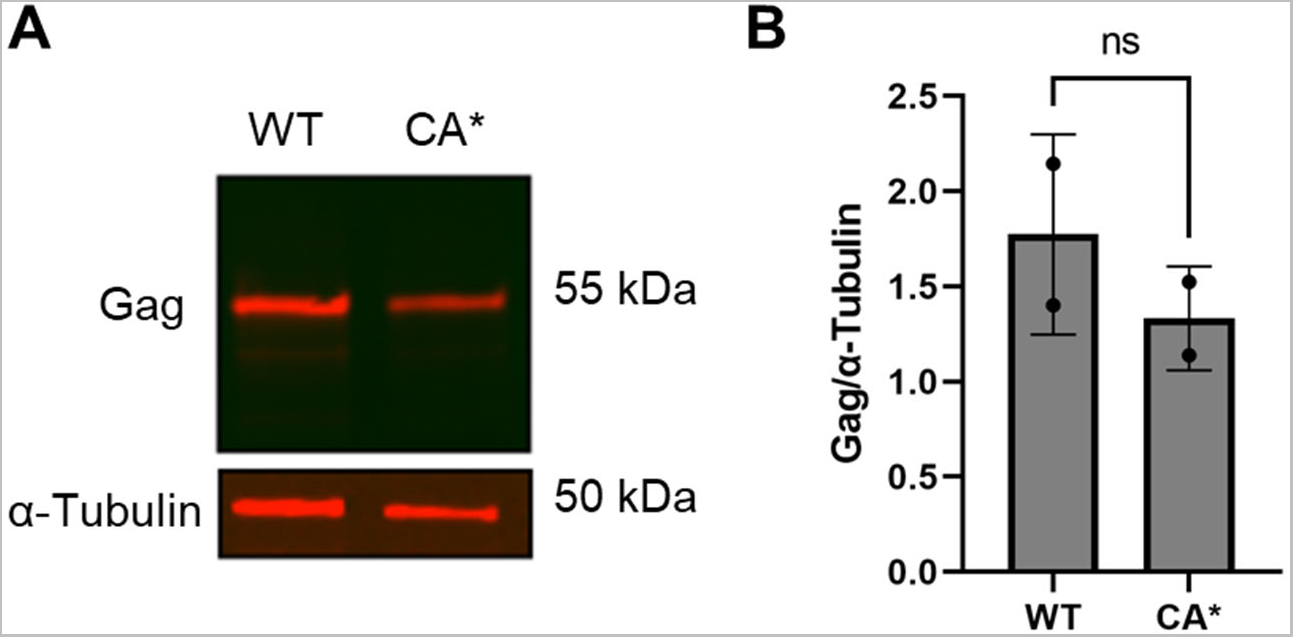
Estimation of CA* and WT CA stoichiometry upon virus production. **A)** Fluorescent western blot showing SQV (200 nM) treated CA* and WT Gag from HEK293T/17 cells. CA* and WT Gag was detected using the human anti-HIV serum antibody and visualized with goat anti-human 800CW secondary antibody. GAPDH loading control is below the Gag western blot. GAPDH was detected using rabbit anti-α-tubulin antibodies visualized with Goat anti-rabbit 800CW secondary antibody. **B)** Densitometry analysis of Gag/tubulin ratio is shown. Two-independent experiments with mean and S.D. plotted.

**Fig. S3.**
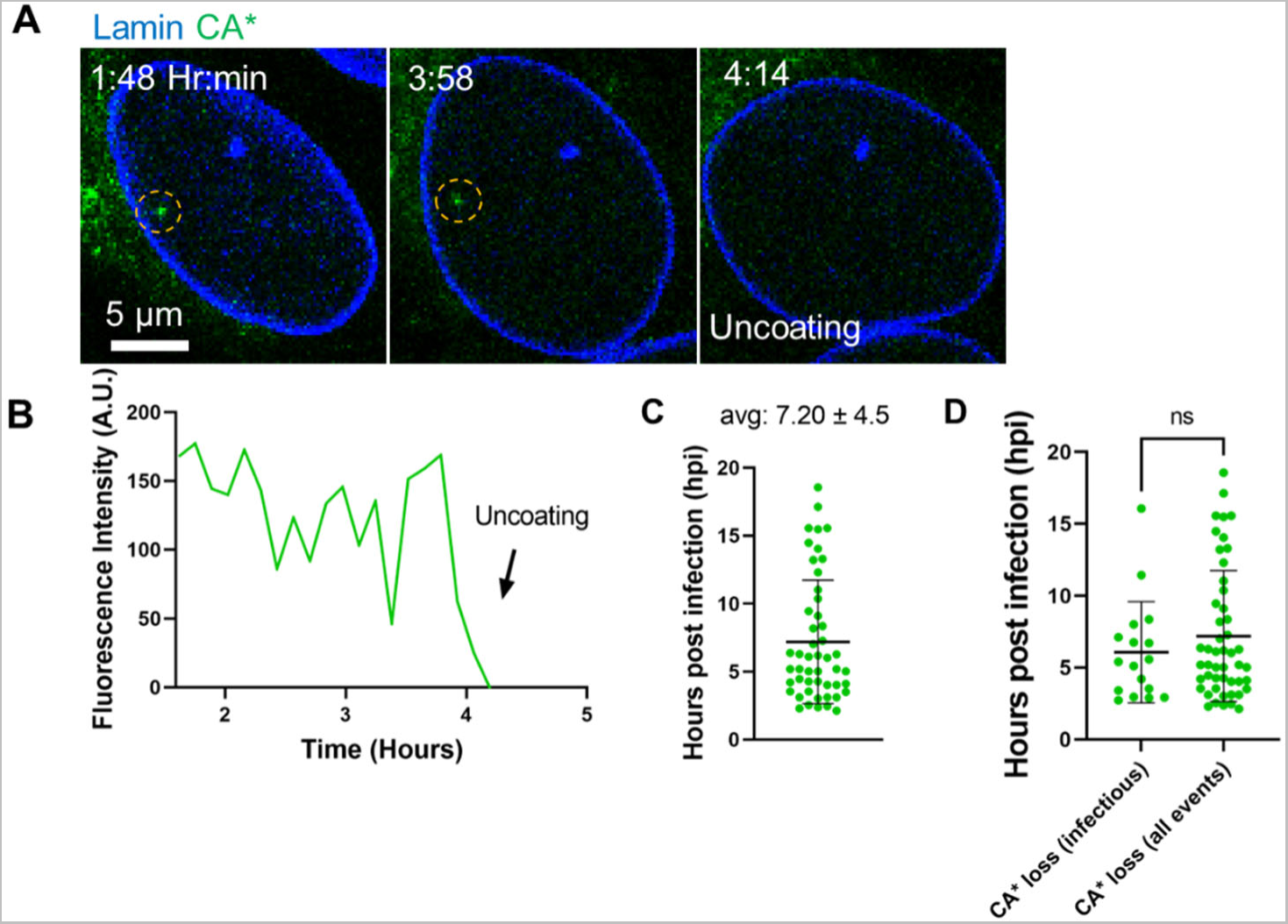
Kinetics of nuclear uncoating events that are imaged without monitoring infection. **A)** Representative micrograph of CA* uncoating in the nucleus of a GHOST-SNAP-lamin cell. Yellow dashed circle highlights the location of the CA* core of interest. **B)** Single particle intensity trace of the nuclear CA* core from panel A. The uncoating event (terminal capsid loss) is marked by an arrow. **C)** Kinetics of nuclear uncoating events of GHOST-SNAP-lamin cells (n = 49). Average uncoating times with standard deviations are shown above the columns. **D)** Infectious uncoating events from Fig. 3C were compared to all uncoating events that were imaged without GFP reporter expression from Fig. S3C. No significant difference was visualized in the uncoating kinetics between infectious and all uncoating events (p = 0.54, Mann-Whitney rank sum test).

**Fig. S4.**
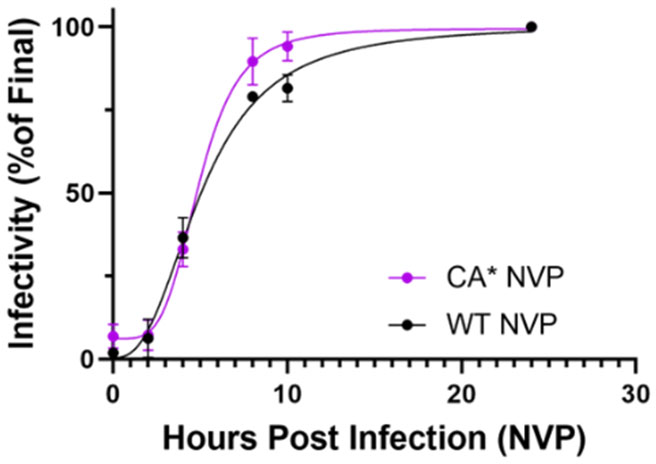
NVP time-of-addition assay to assess reverse transcription kinetics. Representative reverse transcription kinetics of CA^WT^ and CA* pseudoviruses treated with 10 µM NVP at 0, 2, 4, 8, 10, 24 hpi. The T_50_ for WT and CA* pseudoviruses are 5.1 and 4.9 hpi, respectively. Mean and S.D. plotted between 2 independent virus preparations.

**Fig. S5.**
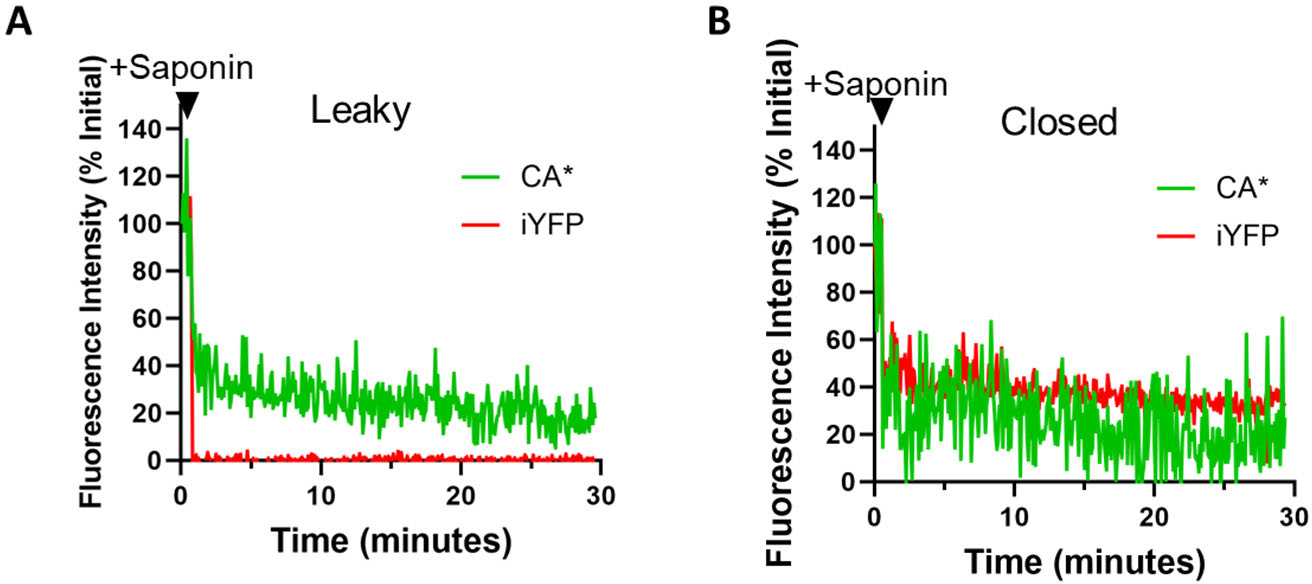
Single particle intensity traces for leaky and closed CA*:Gag-iYFP cores *in vitro*. **A)** single particle intensity trace for coverglass-immobilized CA*:Gag-iYFP pseudovirus displaying the leaky iYFP loss phenotype. **B)** Single particle intensity trace for CA*:Gag-iYFP pseudovirus displaying the closed phenotype, never losing iYFP single from mature core.

**Fig. S6.**
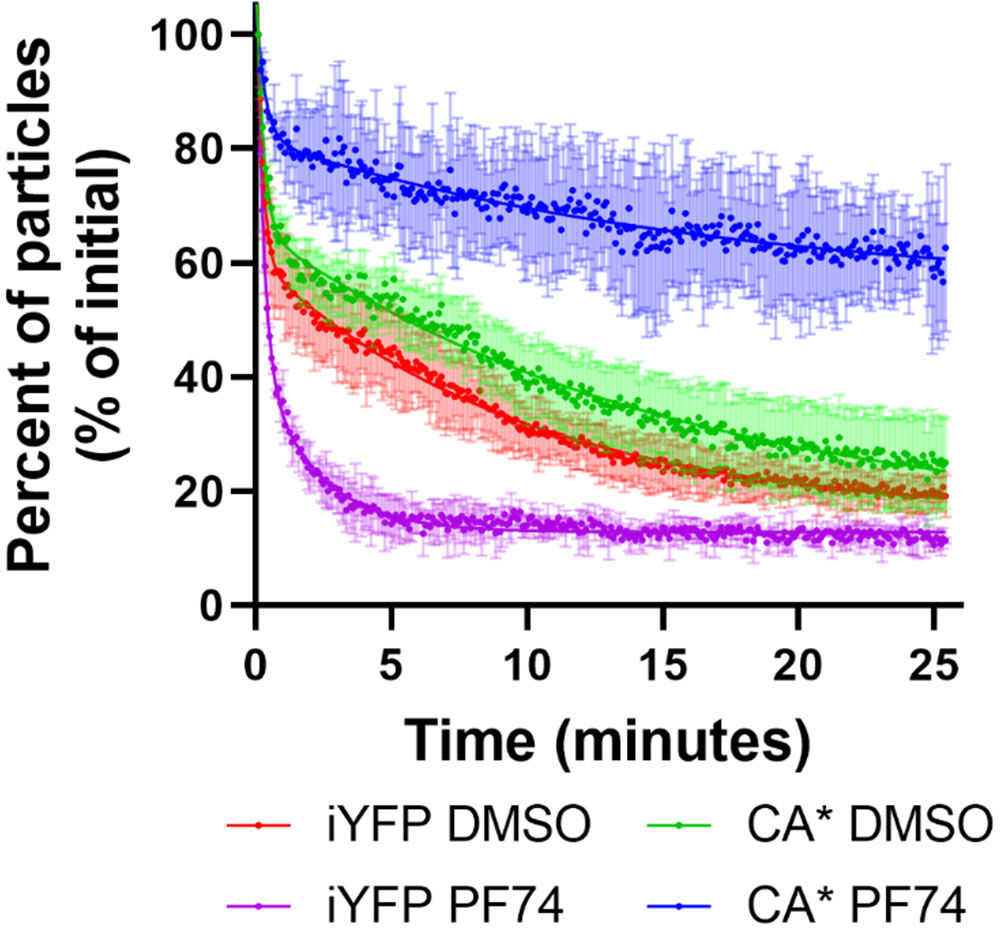
Release of core-trapped iYFP from in vitro CA*:Gag-iYFP cores. Kinetics of uncoating (reduction in number of particles per image field) for CA*:Gag-iYFP cores *in vitro,* following saponin treatment (100 µg/mL, t=0) in the presence of 10 µM PF74 or DMSO. Particle counts were normalized to the initial quantity. Immature particles were excluded from analysis. Non-zero plateaus are the result of cores that retain above-threshold amounts of iYFP or CA*. Means and S.D. are from 2 independent experiments.

**Fig. S7.**
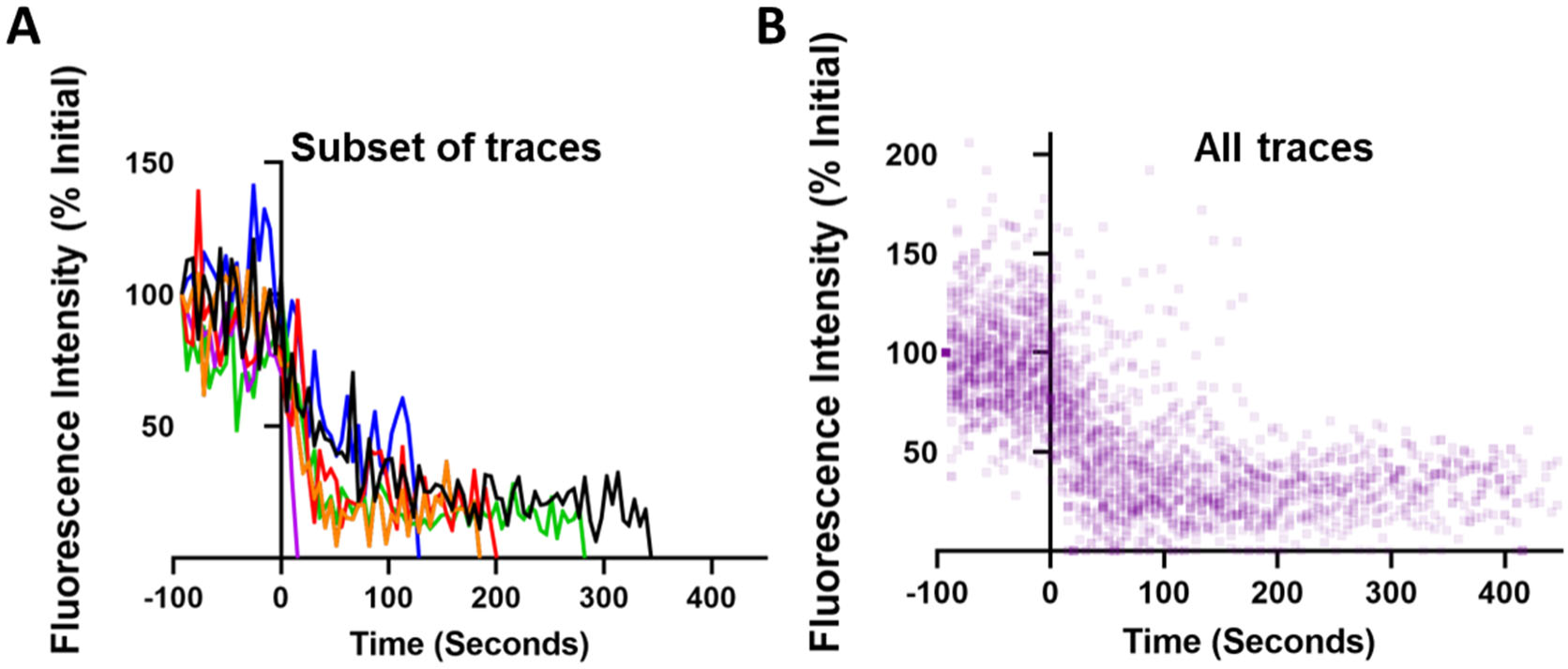
Single particle intensity traces of all *in vitro* integrity loss and uncoating events from Fig. 4J. **A)** Single particle intensity traces from the ensemble plot in Fig, 4F. A small subset of traces was selected to demonstrate phenotypes and have an individual color. **B)** All traces were selected, and each time point is represented with a transparent point to demonstrate areas of overlap. Most particles undergo near-immediate loss of CA* fluorescence after integrity loss.

**Table S1. Primers designed for cloning the CA* viral vectors.** PCR primers with cloning restriction sites in violet, and mutagenesis sites in red.

## Supplementary Movie Legends

**Movie S1. Live cell imaging of CA* nuclear import.** Representative movie of a single CA* core entering the nucleus of a GHOST-SNAP-Lamin cell at early times post infection without losing CA* fluorescence. SNAP-Lamin B1 is labeled with SNAP-OregonGreen. Red arrows point to cores of interest with annotations signifying the nuclear import event.

**Movie S2. HIV-1 nuclear uncoating is manifested by a terminal loss of CA* signal.** Representative movie demonstrating terminal loss of the CA* signal within the nucleus of a GHOST-SNAP-Lamin cell. SNAP-Lamin B1 is labeled with SNAP-OregonGreen. Red arrows point to CA* core of interest with annotation demonstrating the uncoating process.

**Movie S3. Nuclear uncoating correlates with productive infection.** Representative movie for terminal nuclear uncoating of a single CA* core (green) in the nucleus of a GHOST-SNAP-Lamin cell (MOI 0.1). SNAP-Lamin B1 is labeled with SNAP-TMR-STAR. Uncoating and GFP reporter expression (red) are annotated, and red arrow points to core of interest.

**Movie S4. PF74 leads to release of core-trapped fluid phase marker and results in capsid degradation.** Representative movie showing PF74-induced loss of integrity (iYFP loss, red) of a single CA*:Gag-iYFP core followed by degradation of the capsid lattice (loss of CA* signal, green). SNAP-Lamin B1 is labeled with SNAP-TMR-STAR. Arrows point to the CA*:Gag-iYFP core of interest with annotations signifying the integrity loss and CA* degradation events.

**Movie S5. Multi-step nuclear HIV-1 uncoating in a GHOST-SNAP-Lamin cell.** Representative movie of a single CA*:Gag-iYFP labeled core that loses of integrity (releases iYFP, red) and uncoats (loses CA*, green) within the nucleus of a GHOST-SNAP-Lamin cell. SNAP-Lamin B1 is labeled with SNAP-TMR-STAR. Arrows point to CA*:Gag-iYFP core of interest with annotations signifying the integrity loss and uncoating events.

**Movie S6. Multi-step uncoating in the nucleus of THP-1 macrophage-like cells.** Representative movie of a single CA*:Gag-iYFP core undergoing loss of integrity (release of iYFP, red) followed by terminal uncoating (loss of CA*, green) within the nucleus in a differentiated THP-1 macrophage cell. Nucleus is stained with Hoechst 33342 (blue). Arrows point to CA*:Gag-iYFP core of interest with annotations signifying the integrity loss and uncoating events.

